# A TARGETED COMBINATION THERAPY ACHIEVES EFFECTIVE PANCREATIC CANCER REGRESSION AND PREVENTS TUMOR RESISTANCE

**DOI:** 10.1101/2025.08.04.668325

**Authors:** Vasiliki Liaki, Sara Barrambana, Myrto Kostopoulou, Carmen G. Lechuga, Ruth Álvarez, Rebeca Barrero, Silvia Jiménez-Parrado, Alejandra López-García, Lucia Morales-Cacho, Juan Carlos López-Gil, Elena Zamorano-Dominguez, Pian Sun, Blanca Rosas-Perez, Marta San Roman, Eduardo Caleiras, Matthias Drosten, Bruno Sainz, Monica Musteanu, Nelson Dusetti, Valeria Poli, Francisco Sánchez-Bueno, Carmen Guerra, Mariano Barbacid

**Author notes:** Co-senior authors (Carmen Guerra, Mariano Barbacid). Correspondence (C.G.), (M.B.).

## Abstract

Pancreatic ductal adenocarcinoma (PDAC) has one of the lowest cancer survival rates. Recent studies using RAS(ON) inhibitors as single agents have opened the door to more efficacious therapies. Here, we demonstrate that genetic ablation of three independent nodes involved in downstream (RAF1), upstream (EGFR) and orthogonal (STAT3) KRAS signaling pathways leads to complete and permanent disappearance of orthotopic PDACs induced by KRAS/TP53 mutations. Likewise, a combination of RAS(ON) (RMC-6236/daraxonrasib), EGFR family (afatinib) and STAT3 (SD36) selective inhibitors/degraders induced the effective regression of these orthotopic tumors with no evidence of tumor resistance for over 200 days post-treatment. This combination therapy also led to significant regression of genetically engineered mouse tumors as well patient-derived tumor xenografts (PDX) in the absence of tumor relapses. Finally, this combination therapy was well tolerated by the animals. These results should guide the development of clinical trials that could benefit PDAC patients.

## INTRODUCTION

Pancreatic ductal adenocarcinoma (PDAC) has become the third leading cause of carcer related deaths in the Western World, only second to colorectal and lung cancer^1^. These high levels of mortality are mainly attributed to the lack of efficacious therapies. This situation is paradoxical, considering the significant progress made during the last two decades on the biology of the disease, mainly thanks to the development of genetically engineered mouse models that faithfully reproduce the natural history of the human disease^2,3^. Current therapeutic options available to PDAC patients still rely on cytotoxic drugs such as gemcitabine, approved almost thirty years ago^4^, either alone or in combination with nab-paclitaxel. Other highly toxic drug combinations such as FOLFIRINOX offer limited benefit and can only be used in fit patients^5^. KRAS oncogenes, the most common initiating event, were considered undruggable until recently^6^. Efforts to target their canonical signaling pathways such as the MAPK and the PI3K cascades have proved to be ineffective in clinical trials due to their high toxicities. This situation took an unexpected turn of events when Shokat and colleagues identified a small pocket capable of housing selective inhibitors^7^. This discovery led to the development of selective drugs targeting the KRAS^G12C^ isoform, such as sotorasib and adagrasib, which have recently gained FDA approval^8-10^. Emerging results using these inhibitors in pancreatic tumors carrying KRAS^G12C^ mutations have resulted in progression free and overall survival of just 4 and 7 months, respectively^11^. New RAS(ON) inhibitors capable of blocking all or most oncogenic KRAS isoforms as well as their normal counterparts have also been recently reported^12-16^. In early clinical data, these inhibitors appear to be more effective^16,17^. In experimental mouse tumor models, these RAS(ON) inhibitors have led to increase in survival. Yet, mice ended up succumbing to the disease due to the appearance of tumor resistance ^14-16^. Hence, there is still an urgent need to develop novel and more efficacious therapeutic strategies against KRAS driven pancreatic cancer, most likely drug combinations that prevent or at least thwart tumor resistance^18-20^.

Previous studies in our laboratory have illustrated that concomitant ablation of RAF1 and EGFR, two mediators of KRAS signaling, induced complete regression of a limited subset of mouse PDAC tumors^21^. We now report that targeting an additional signaling node, STAT3, in combination with RAF1 and EGFR, led to the complete and durable regression of mouse and human PDAC tumors. More importantly, we have extended these observations to a pharmacological setting by demonstrating that combined inhibition of KRAS, EGFR and STAT3 induces significant and long-lasting regression of these experimental PDACs without inducing significant toxicities.

## RESULTS

### Differential tumor responses to RAF1 and EGFR ablation

*Kras*^+/FSFG12V^;*P53*^F/F^;Elas-tTA;TetO-Flpo;*Rosa26*CreERT2 mice (KPeFC mice) develop PDAC tumors upon expression of the Flpo recombinase in acinar cells during the late stages of embryonic development^21^. Addition of the *Raf*1^L/L^ and *Egfr*^L/L^ floxed alleles allowed us to identify two differential tumor phenotypes based on their response to *Raf*1 and *Egfr* ablation induced upon tamoxifen (TMX) exposure^21^. We have now extended these observations to illustrate a direct correlation between tumor size and response to *Raf*1 and *Egfr* ablation *in vivo*, with tumors larger than 100 mm^3^ being consistently resistant to this therapeutic strategy (Figure 1A). The differential response to *Raf*1 and *Egfr* ablation was also observed when tumors were explanted in culture and these targets eliminated *in vitro*. Of 61 cell lines generated from KPeFC;*Raf1*^L/L^;*Egfr*^L/L^ tumors, 19 completely stopped proliferating upon recombination of the *Raf1* and *Egfr* floxed alleles, whereas 22 cell lines were unaffected. Interestingly, these 22 cell lines came from the biggest tumors (>100 mm^3^). *Raf1* and *Egfr* resistant cell lines continued proliferating at the same pace as those expressing the targets. Noteworthy, 20 tumor cell lines exhibited a mixed phenotype with different proportions of cells surviving upon *Raf1* and *Egfr* ablation (Figure 1B,C and Figure S1A). Subclones derived from the “Mixed” cell lines no longer displayed a mixed phenotype. Instead, they behave as *bona fide* sensitive or resistant cells (Figure 1D,C). Finally, sensitive tumor cells acquired a resistant phenotype after implantation in immunocompetent or immunodeficient mice but not upon extensive passage in culture (Figure 1B,C). These results suggest that pancreatic tumor cells evolve from a sensitive to a resistant phenotype during tumor progression *in vivo*. RNA seq analysis of pre- and post-implanted tumor cells revealed enriched signaling pathways related to immune and stromal microenvironment (Figure S1D). Thus, adding further support to the role of the tumor microenvironment in this process.

**Figure 1.**
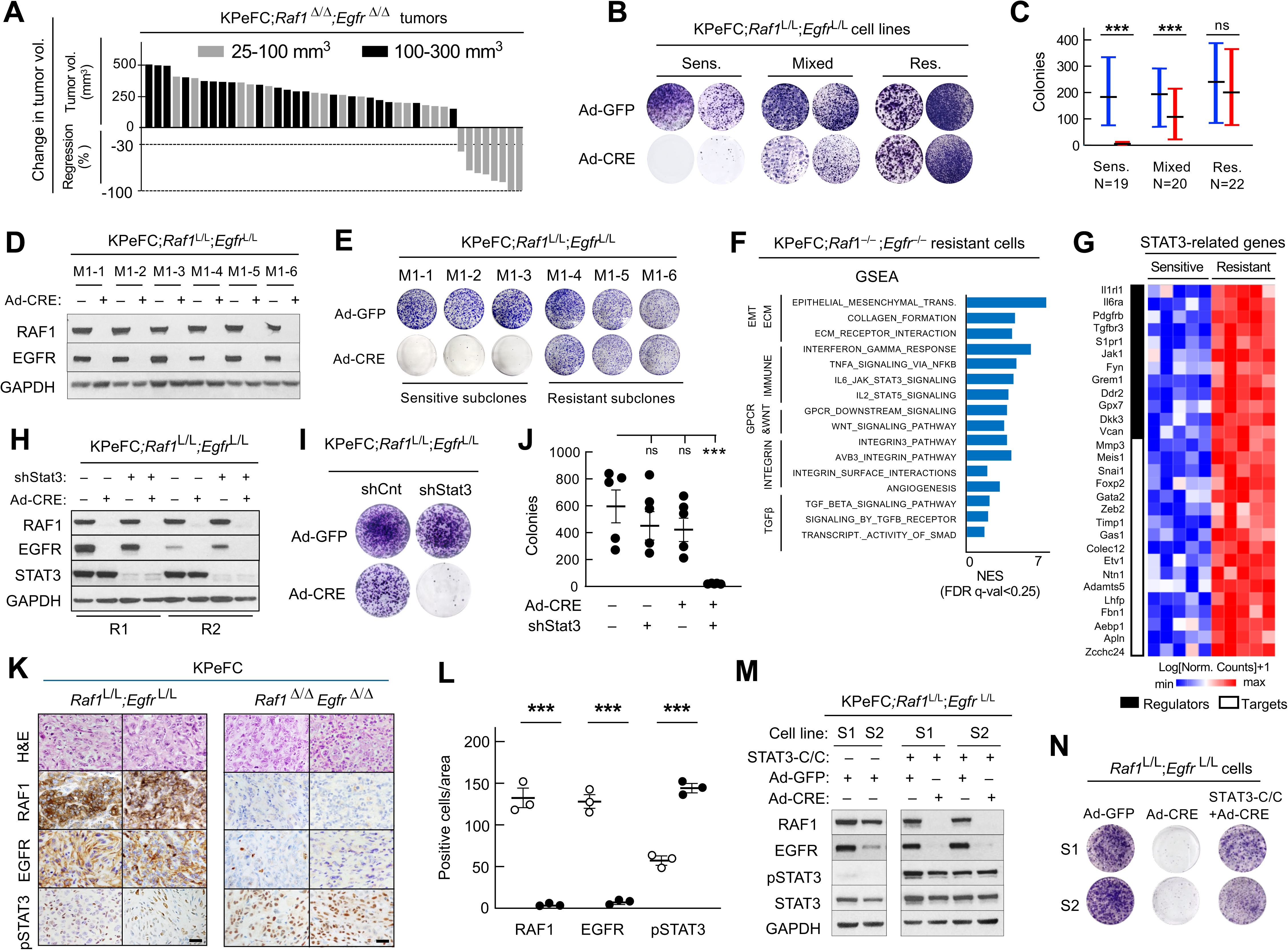
Differential response of mouse PDAC tumors to *Raf1* and *Egfr* ablation. **A,** Waterfall plot representing changes in tumor volume of tumors present in KPeFC;*Raf1*^L/L^;*Egfr*^L/L^;*Rosa26*CreERT2 mice exposed to a TMX diet as determined by ultrasound. Tumors ranging in size from 25 to 100 mm^3^ at the start of the treatment are represented by gray columns. Tumors of sizes ranging from 100 to 300 mm^3^ at the start of the treatment are represented by solid columns. **B,** Colony formation assay of representative KPeFC;*Raf1*^L/L^;*Egfr*^L/L^ tumor cell lines classified as Sensitive, Mixed or Resistant based on their response to *Raf*1 and *Egfr* ablation. **C,** Quantification analysis of the number of colonies displayed by Sensitive, Mixed and Resistant cell lines before (blue columns) and after (red columns) ablation of *Raf1*^L/L^ and *Egfr*^L/L^ conditional alleles indicate the range of the values. Solid horizontal bars indicate the mean values. Numbers indicate the number of cells lines in each category. **D,** Western blot analysis of RAF1 and EGFR expression in whole cell extracts of subclones derived from a representative “Mixed” tumor cell line (M1-1 to M1-6) infected with Adeno-GFP (–) or Adeno-CRE (+) viral particles. GAPDH was used as loading control. **E,** Phenotypic identification of these subclones as Sensitive or Resistant to *Raf*1 and *Egfr* ablation by colony assay after infection with Adeno-GFP or Adeno-CRE viral particles. **F,** GSEA analysis of pathways of Resistant tumor cell lines after *Raf*1 and *Egfr* ablation. The normalized enrichment score (NES) ranking was generated by GSEA. Selective pathways are represented with q-FDR value <0.25. **G,** Heatmap of the expression levels (Log_2_-transformed values of count+1) of STAT3-related genes in RAF1/EGFR Sensitive (n=5) and Resistant (n=5) tumor cell lines. Those genes that serve as regulators of STAT3 activity are indicated by a solid bar. Those that are known to serve as targets for STAT3 are indicated by an open bar. **H,** Western blot analysis of EGFR, RAF1 and STAT3 expression levels in whole cell-extracts of two independent RAF1/EGFR Resistant tumor cell lines (R1 and R2) expressing control shRNAs or shRNAs against STAT3 followed by infection with Adeno-GFP or Adeno-CRE viral particles. GAPDH served as loading control. **I,** Colony assay of a resistant KPeFC*;Raf1*^L/L^*;Egfr*^L/L^ tumor cell line incubated with control shRNAs or with shRNAs specific for *Stat*3 and infected with Adeno-CRE viral particles to eliminate RAF1 and EGFR expression. **J**, Quantification of the number of colonies shown in **I.** The P value was obtained using paired two-tailed t test. ns, not significant, *** P<0.001. Error bars indicate mean ± SEM. **K,** H&E and IHC analysis of RAF1, EGFR, and pSTAT3^Y705^ expression in sections of representative (**left**) RAF1/EGFR Resistant KPeFC;*Raf*1^L/L^;*Egfr*^L/L^ (n=2) or (**right**) KPeFC;*Raf1*^Δ/Δ^;*Egfr* ^Δ/Δ^ (n=2) tumors. Scale bar represents 50 μm. **L**, Quantification of the number of positively stained cells in KPeFC;*Raf*1^L/L^;*Egfr*^L/L^ tumors (open circles) and in KPeFC;*Raf1*^Δ/Δ^;*Egfr* ^Δ/Δ^ tumors (solid circles) shown in **K.** The P value was obtained using paired two-tailed t test, *** P<0.001. Error bars indicate mean ± SEM**. M,** Western blot analysis of the expression of RAF1, EGFR, pSTAT3 and STAT3 in cell extracts of two independent Sensitive KPeFC*;Raf1*^L/L^*;Egfr*^L/L^ tumor cell lines, S1 and S2, infected with a cDNA encoding the constitutively active human STAT3-C/C isoform. Cells were infected with Adeno-GFP or Adeno-CRE viral particles, as indicated. **N**, Colony assay of Sensitive KPeFC*;Raf1*^L/L^*;Egfr*^L/L^ S1 and S2 tumor cell lines expressing the human STAT3-C/C isoform and infected with Adeno-GFP or Adeno-CRE viral particles.

### Molecular profile of pancreatic tumors sensitive and resistant to *Raf1* and *Egfr* ablation

Principal Component Analysis (PCA) of transcriptomic data of 13 sensitive and 14 resistant tumor cell lines confirmed that they have distinct molecular profiles with a significant number of differentially expressed genes (Figure S1E and Supplementary Table 1) as previously described^21^. Moreover, Gene Set Enrichment Analysis (GSEA) showed increased expression of KRAS, ERBB, PI3K and WNT signaling in sensitive tumor cells, along with other pathways such as fatty and bile acid metabolism that have been associated with the classical/progenitor subtypes defined for human PDACs (Figure S1F). On the other hand, resistant tumor cells displayed enriched EMT extracellular matrix signatures linked to the basal/squamous PDAC subtypes (Figure S1G)^22,23^. Tumor cells resistant to the elimination of RAF1 and EGFR expression were also characterized by increased angiogenesis, inflammatory and interferon responses, as well as signaling mediated by STAT3 and STAT5, NFkB and SMAD (Figure S1G).

### Molecular profile of resistant tumor cells devoid of RAF1 and EGFR expression

To interrogate the mechanisms responsible for the resistance of PDAC tumor cells to *Raf1* and *Egfr* ablation, we submitted 14 independent RAF1/EGFR resistant tumor cell lines to RNA seq analysis before and after elimination of these genes. GSEA analysis revealed upregulation of gene sets related to EMT, collagen formation and immune pathways, including interferon response, JAK-STAT and NFkB signaling in the ablated cells (Figure 1F). We also detected increased Integrin- and GPCR-mediated signaling, as well as angiogenesis and TGFβ signaling (Figure 1F). To validate the potential therapeutic value of these upregulated pathways, we targeted the expression of NFkB, TGFβ and GPCR signaling using shRNAs against the p65 *RelA* subunit of the NFkB complex and against the genes encoding TGFBR3 and S1PR1, two receptors involved in the TGFβ and GPCR pathways, respectively (Figure S1H). As illustrated in Figure S1I, their inhibition had no effect on the proliferation of resistant tumor cells. Similar results were observed when these tumor cells were treated with the TGFβ inhibitor, galunisertib (Figure S1J).

### RAF1/EGFR resistant tumor cells undergo apoptotic cell death upon silencing of STAT3 expression

Transcriptomic analysis also revealed activation of the STAT3 signaling pathway, including 12 genes involved in the regulation of STAT3 activity as well as 17 STAT3 target genes (Figure 1F,G). These observations reinforced previous results indicating that STAT3 became activated by phosphorylation on Tyr^705^ upon *Raf1* and *Egfr* ablation^21^. To determine whether the activation of STAT3 had any functional consequences on the proliferating properties of the resistant tumor cells, we targeted *Stat*3 expression with specific shRNAs before and after elimination of *Raf1* and *Egfr* (Figure 1H). *Stat*3 silencing led to the rapid death of the resistant cells in the absence of RAF1 and EGFR expression (Figure 1I,J). Moreover, we observed increased expression of pSTAT3^Y705^ in KPeFC;*Raf1*^-/-^;*Egfr*^-/-^ PDAC tumors, compared with tumors expressing RAF1 and EGFR (Figure 1K,L). To further illustrate the role of STAT3 in the resistance of PDAC tumor cells to RAF1/EGFR ablation, we performed rescue experiments *in vitro* using a cDNA encoding a constitutively active isoform of human STAT3 (STAT3^A661C/N663C^), designated as STAT3-C/C (ref. 24). Infection of three independent KPeFC;*Raf1*^L/L^;*Egfr*^L/L^ tumor cell lines sensitive to RAF1/EGFR ablation with a vector expressing the STAT3-C/C isoform abrogated their sensitivity to RAF1/EGFR ablation and induced proliferation of these cells after removal of RAF1/EGFR expression (Figure 1M,N).

### Apoptotic tumor cell death upon concomitant ablation of *Raf1, Egfr* and *Stat3* loci

To interrogate the effect of ablating *Raf1, Egfr* and *Stat3*, we generated a mouse strain carrying conditional alleles for the three targets, KPeFC;*Raf1*^L/L^;*Egfr*^L/L^;*Stat3*^L/L^ (ref. 21,25). Tumor cells were obtained from 30 independent tumors. Cells resistant to RAF1/EGFR ablation were identified by dual silencing of RAF1 and EGFR expression with specific shRNAs (Figure S2A,B). Adeno-CRE infection of these resistant KPeFC;*Raf1*^L/L^;*Egfr*^L/L^;*Stat3*^L/L^ tumor cell lines consistently resulted in cell death upon elimination of the three loci (Figure 2A,B). However, elimination of only two loci, in all possible combinations, had no effect on tumor cells proliferation (Figure 2A). Hence, expression of just one of the three signaling nodes, either RAF1, EGFR or STAT3, was sufficient to maintain tumor cell proliferation.

**Figure 2.**
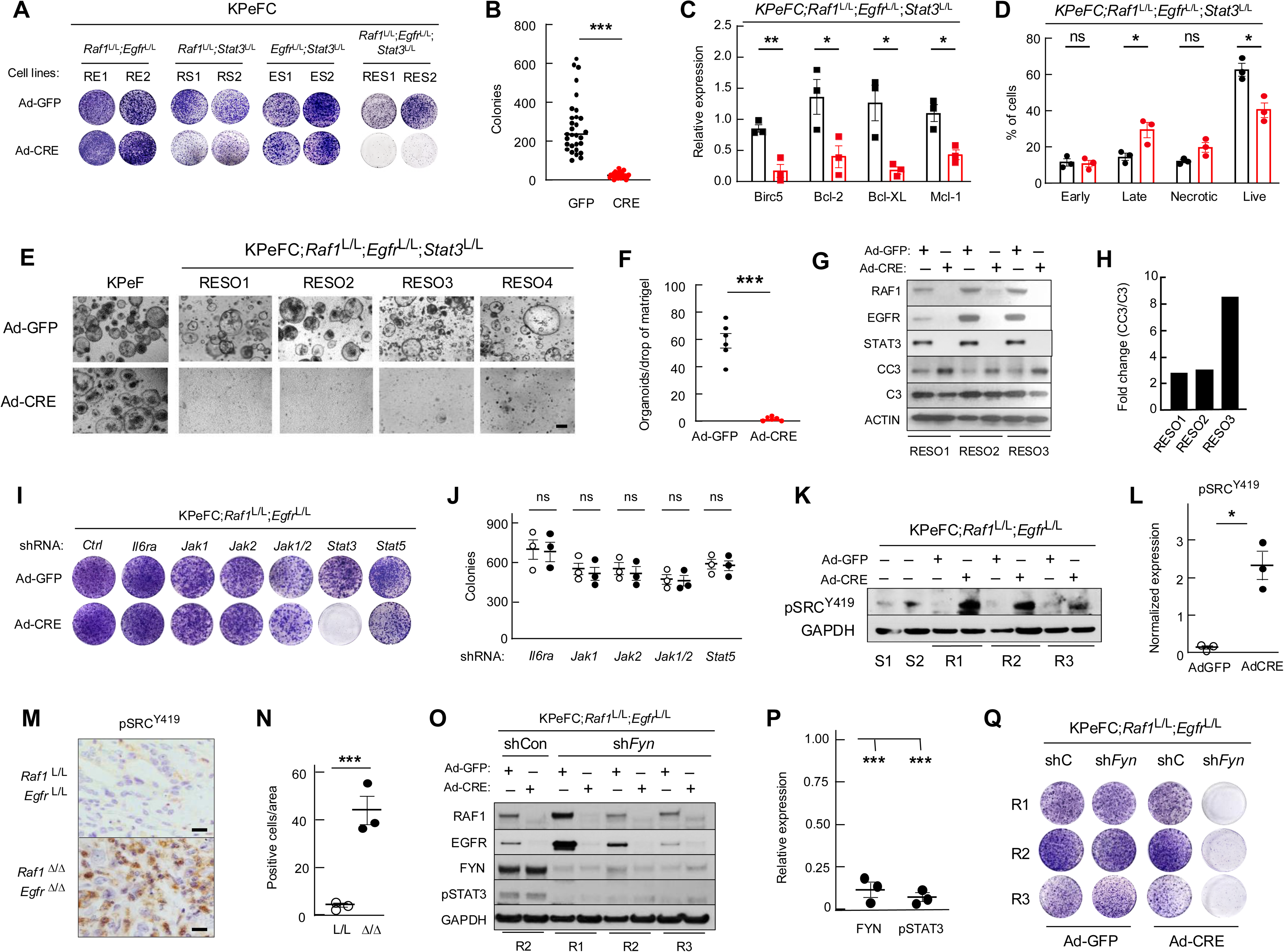
STAT3 mediates tumor resistance to *Raf*1 and *Egfr* ablation. **A,** Colony assay of two representative Resistant KPeFC*;Raf1*^L/L^*;Egfr*^L/L^ (RE1, RE2) KPeFC*;Raf1*^L/L^*;Stat3*^L/L^ (RS1, RS2), KPeFC*;Egfr*^L/L^*;Stat3*^L/L^ (ES1, ES2) and KPeFC*; Raf1*^L/L^*;Egfr*^L/L^*;Stat3*^L/L^ (RES1, RES2) pancreatic tumor cell lines infected with Adeno-GFP or Adeno-CRE viral particles, as indicated. **B,** Quantification of the number of colonies of KPeFC*;Raf1*^L/L^*;Egfr*^L/L^: *Stat3*^L/L^ tumor cells (n=30) infected with Adeno-GFP (solid squares) or Adeno-CRE (red circles) viral particles. The P value was obtained using paired two-tailed t test. *** P<0.001. Error bars indicate mean ± SEM. **C**, Expression levels of apoptosis related genes including Bicr5, Bcl-2, Bcl-XL and Mcl-1 determined by qRT-PCR in three independent KPeFC*;Raf1*^L/L^*;Egfr*^L/L^;*Stat3*^L/L^ tumor cell lines, three days after infection with Adeno-GFP (solid squares) or Adeno-CRE (red squares) viral particles. Open bars represent the mean values. The P value was obtained using paired two-tailed t test. * P<0.05, ** P<0.01. Error bars indicate mean ± SEM. **D**, Percentage of early and late apoptotic, necrotic and live cells of representative (n=3) Resistant KPeFC*;Raf1*^L/L^*;Egfr*^L/L^: *Stat3*^L/L^ tumor cell lines as determined by flow cytometry analysis after Annexin V and DAPI staining, three days after infection with Adeno-GFP (solid circles) or Adeno-CRE (red circles) viral particles. Open columns represent the mean values. The P value was obtained using paired two-tailed t test, ns, not significant, * P<0.05. Error bars indicate mean ± SEM. **E**, Representative mouse organoid cultures established from individual RAF1/EGFR Resistant tumors (RESO1 to RESO4) of KPeFC;*Raf1*^L/L^;*Egfr*^L/L^;S*tat3*^L/L^ mice 7 days after infection with Adeno-GFP or Adeno-CRE viral particles. An organoid culture derived from a tumor isolated from a KPeF mouse was used as control. Images were obtained with a confocal microscope. Scale bar represents 200 μm. **F,** Quantification of the number of organoids per drop of matrigel of six independent organoids derived from KPeFC*;Raf1*^L/L^*;Egfr*^L/L^;*Stat3*^L/L^ tumors infected with Adeno-GFP (solid circles) or Adeno-CRE (red circles) viral particles. The P value was obtained using paired two-tailed t test. *** P<0.001. Error bars indicate mean ± SEM. **G**, Western blot analysis of RAF1, EGFR, STAT3, CC3 and C3 expression in cell extracts derived from three independent KPeFC*;Raf1*^L/L^*;Egfr*^L/L^: *Stat3*^L/L^ tumor organoids (RESO1 to RESO3) three days after infection with Adeno-GFP or Adeno-CRE viral particles, as indicated. Actin served as loading control. **H**, Fold change (Adeno-CRE vs. Adeno-GFP) of the ratio of CC3 and C3 expression normalized with Actin as indicated in **G**. **I**, Colony formation assay of a representative RAF1/EGFR Resistant KPeFC;*Raf1*^L/L^;*Egfr*^L/L^ tumor cell line expressing the indicated shRNAs. Cells were infected with Adeno-GFP or Adeno-CRE viral particles. **J**, Quantification of the number of colonies of three independent Resistant KPeFC;*Raf1*^L/L^;*Egfr*^L/L^ tumor cell lines as shown in **I**. **K,** Western blot analysis of phosphorylated members of the SRC kinase family using a pSRC^Y419^ antibody in whole cell-extracts of two independent RAF1/EGFR Sensitive (S1 and S2) and three Resistant (R1 to R3) tumor cell lines infected with Adeno-GFP or Adeno-CRE viral particles. GAPDH was used as loading control. **L**, Quantification of the relative expression of pSRC^Y419^ normalized to the loading control shown in **K** infected with Adeno-GFP (open circles) or Adeno-CRE (solid circles) viral particles. The P value was obtained using paired two-tailed t test. * P<0.05. Error bars indicate mean ± SEM. **M,** IHC analysis of the expression of phosphorylated members of the SRC kinase family using a pSRC^Y419^ antibody in sections of tumors obtained from a representative RAF1/EGFR Resistant KPeFC;*Raf1*^L/L^;*Egfr*^L/L^ and KPeFC;*Raf1* ^Δ/Δ^;*Egfr* ^Δ/Δ^ mouse. Scale bar represents 50 μm. **N**, Quantification of the number of positive pSRC^Y419^ cells in KPeFC;*Raf1*^L/L^;*Egfr*^L/L^ (open circles) and KPeFC;*Raf1* ^Δ/Δ^;*Egfr* ^Δ/Δ^ (closed circles) tumors, (n=3). The P value was obtained using unpaired two-tailed t test. *** P<0.001. Error bars indicate mean ± SEM. **O**, Western blot analysis of RAF1, EGFR, FYN and pSTAT3 expression in whole cell-extracts of three representative RAF1/EGFR Resistant KPeFC;*Raf1*^L/L^;*Egfr*^L/L^ tumor cell lines (R1 to R3) expressing control shRNAs (shCon) or shRNAs against *Fyn* (sh*Fyn*) after infection with Adeno-GFP or Adeno-CRE viral particles. GAPDH was used as loading control. **P**, Quantification of the relative expression levels of FYN and pSTAT3 in the Western blot shown in **O** in the presence of shFyn respect to those in the presence of shControl (shCon). The P value was obtained using unpaired two-tailed t test. *** P<0.001. Error bars indicate mean ± SEM. **Q,** Colony formation assay of three representative RAF1/EGFR Resistant KPeFC;*Raf1*^L/L^;*Egfr*^L/L^ tumor cell lines (R1 to R3) expressing control shRNAs (shC) or shRNAs against *Fyn* (sh*Fyn*). Cells were infected with Adeno-GFP or Adeno-CRE particles as indicated.

As illustrated in Figure 2C, we observed three days after adeno-CRE infection a significant downregulation of apoptotic genes including *Birc5, Bcl-2, Bcl-XL* and *Mcl-1.* In addition, cytometric analysis with antibodies against Annexin-V revealed a significant reduced proportion of live cells along with an increase number of cells in late apoptosis (Figure 2D). Similar results were observed using organoids derived from an independent set of KPeFC;*Raf1*^L/L^;*Egfr*^L/L^;*Stat3*^L/L^ tumors (Figure 2E-H). These results demonstrate that concomitant ablation of *Raf1, Egfr* and *Stat3* induces the elimination of mouse PDAC cells via apoptotic mechanisms. Finally, silencing of S*tat*5, the other member of the STAT family frequently activated in PDAC (ref. 26) had no effect on the proliferation of resistant cells upon ablation of RAF1 and EGFR expression (Figure 2I,J and Figure S2E).

### Mechanism of activation of STAT3

Next, we interrogated the mechanism(s) responsible for the activation of STAT3 via phosphorylation on Tyr^705^ upon ablation of Raf1 and EGFR (ref. 21). First, we assessed the role of canonical STAT3 regulators such as IL6RA, JAK1 and JAK2 (ref. 27). Treatment of resistant tumor cells with and without expression of RAF1 and EGFR with four independent JAK1/2 inhibitors (ruxolitinib, itacitinib, tofacitinib and baricitinib) did not affect cell viability (Figure S2C). In addition, silencing of either *Jak1* or *Jak2* expression with specific shRNAs had no effect on cell proliferation (Figure 2I,J and Figure S2D). Similar results were obtained when we silenced both *Jak1/Jak2* kinases or with specific shRNAs against the IL6 receptor, IL6RA (Figure 2I,J and Figure S2D).

We next explored the role of the SRC family of kinases in the activation of STAT3 via phosphitylation of residue Tyr^705^. Immunoprecipitation analysis of lysates of resistant tumor cells expressing or lacking RAF1/EGFR expression with anti-pSRC^Y419^ only identified phosphorylated SRC proteins in cells lacking RAF1/EGFR expression (Figure 2K,L). Likewise, we detected increased levels of phosphorylated SRC proteins in tissue sections of KPeFC;*Raf1*^-/-^; *Egfr*^-/-^ tumors compared to those expressing both targets (Figure 2M,N). Transcriptomic analysis of members of this gene family revealed increased expression of FYN (ref. 28) in resistant cells (Figure 1G). Indeed, silencing of *Fyn* expression with specific shRNAs inhibited phosphorylation of STAT3 and selectively induced death of resistant tumor cells lacking RAF1 and EGFR expression (Figure 2O-Q).

### Combined ablation of *Raf*1, *Egfr* and *Stat*3 induces the complete and long-lasting regression of orthotopic pancreatic tumors

We next extended these observations to *in vivo* studies. To this end, we used orthotopic pancreatic tumor models, since the systemic elimination of STAT3 in adult mice results in the death of the animals due to intestinal ulcers and peritonitis (Figure S3A,B)^29-31^. Furthermore, we found additional defects within the myeloid population along the accumulation of immune cells in the periphery of liver tissue and increase of the white pulp in the spleen (Figure S3C,D). Twelve independent RAF1/EGFR resistant KPeFC;*Raf1*^L/L^;*Egfr*^L/L^;*Stat3*^L/L^ tumor cell lines (RES1 to RES12) were implanted in the pancreas of immunocompetent C57BL/6 mice in triplicates, and the resulting tumor-bearing mice were separated into two cohorts (Figure 3A,B and Figure S4A,B). These orthotopic tumor models developed a significant desmoplastic stroma similar to that present in endogenous PDACs. In one cohort, tumors were allowed to progress until they reached a volume of 400-500 mm^3^ (Figure 3A and Figure S4A). In the second cohort, tumor-bearing mice were exposed to a TMX diet when they reached 50-100 mm^3^ to allow CreERT2-mediated recombination of the conditional *Raf1*^L^*, Egfr*^L^ and *Stat3*^L^ alleles (Figure 3B and Figure S4B). As illustrated in Figure 3C and Figure S4C, expression of RAF1, EGFR and STAT3 was efficiently eliminated just 5 days after exposure to TMX leading to the rapid appearance of apoptotic tumor cells (Figure 3C,D). Tumors of TMX treated mice rapidly decreased in size leading to their complete disappearance in four to six weeks as determined by ultrasound (Figure 3E and Figure S4B).

**Figure 3.**
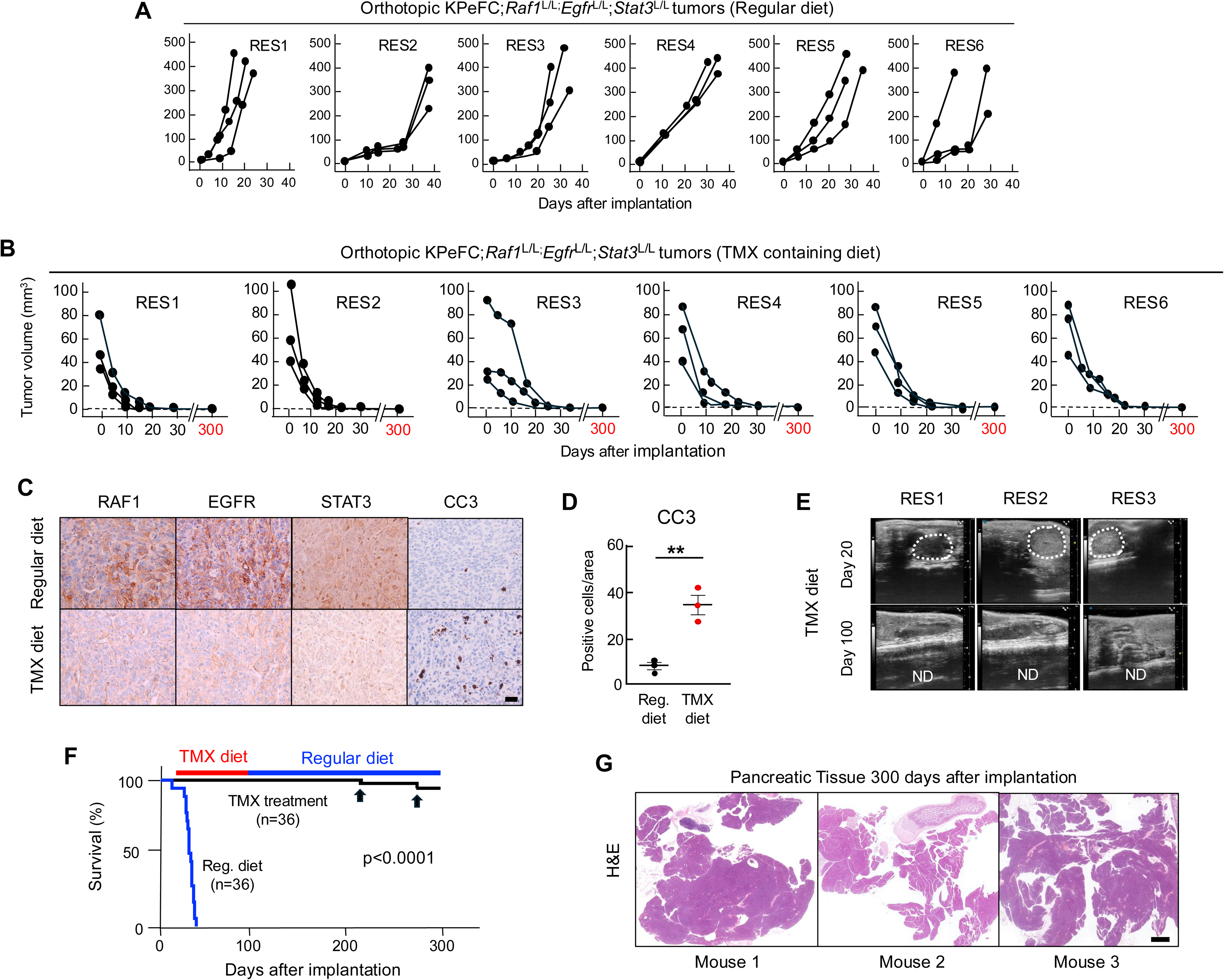
Therapeutic effect of concomitant ablation of *Raf*1, *Egf*r and *Stat3* in orthotopic pancreatic tumors. **A,** Tumor volume visualized by ultrasound imaging of tumors present in C57BL/6 mice (n=3) orthotopically implanted with six RAF1/EGFR resistant KPeFC;*Raf1*^L/L^;*Egfr*^L/L^*;Stat3*^L/L^ tumor cell lines (RES1 to RES6) fed with regular diet (see Figure S4A). **B**, Tumor volume visualized by ultrasound imaging of tumors present in C57BL/6 mice (n=3) orthotopically implanted with six RAF1/EGFR resistant KPeFC; *Raf1*^L/L^;*Egfr*^L/L^*;Stat3*^L/L^ tumor cell lines (RES1 to RES6) exposed to a TMX diet when tumors reached 80 to 100 mm^3^ for 30 days. Once tumors were no longer detected by ultrasound, mice were fed with regular diet until they reached 300 days port-implantation. Regular ultrasound analysis failed to reveale tumor relapse in any of the 18 mice. Dotted line represents tumor sizes undetectable by ultrasound (see Figure S4B). **C**, IHC analysis of RAF1, EGFR, STAT3 and CC3 of tumors of 100 mm^3^ after four days of exposure to either regular or TMX containing diet. Scale bar represents 50 μm. **D**, Quantification of the number of CC3 positive cells per area in depicted in **c**. The results represent the average of CC3 in three independent tumors. The P value was obtained using unpaired two-tailed t test. ** P<0.01. Error bars indicate mean ± SEM. **E**, Representative ultrasound images of the peritoneal cavity of the indicated mice exposed to a TMX diet at the indicated times. Visible tumors are marked with dotted circles. ND, not detected. **F,** Kaplan-Meier survival curve of tumor-bearing mice implanted with 12 independent KPeFC;*Raf1*^L/L^;*Egfr*^L/L^;*Stat3*^L/L^ tumor cell lines (three mice per cell line) and either fed with a regular diet (n=36) (blue line) or exposed to a TMX diet (n=36) for the indicated time (red line). The latter cohort were fed with a regular diet for 200 additional days (straight blue line). Two mice had to be sacrificed at the indicated times (arrows) due to tumor-unrelated causes. The P value was obtained using Log-rank Mantel-Cox test, p<0.0001. **G,** Representative images of H&E-stained sections of pancreatic tissue of mice exposed to the TMX diet for the indicated time and sacrificed at the end of the experiment (300 days post-implantation). No tumoral or stromal tissue could be detected. Scale bar represents 1 mm.

Mice were allowed to thrive without further TMX exposure. None of the mice developed detectable tumors during this time indicating the absence of tumor resistance, one of the main problems of targeted therapies (Figure 3B,F and Figure S4B). Finally, we decided to sacrifice these mice at 300 days after tumor implantation to determine the presence of potential tumor tissue or desmoplastic stroma (Figure 3F). Careful histopathological analysis failed to reveal tumoral or stromal tissue (Figure 3G). These results, taken together, indicate that concomitant ablation of *Raf1, Egfr* and *Stat3* is sufficient to induce the complete regression of advanced PDACs without the appearance of tumor resistance. Moreover, they illustrate that maintenance of the desmoplastic stroma is completely dependent on the presence of tumor cells.

### Combined EGFR and STAT3 inhibitors induce regression of tumors lacking RAF1 expression

To pharmacologically validate this genetic strategy, we first interrogated the effect of EGFR and STAT3 inhibitors in the absence of RAF1 expression since there are no selective RAF1 inhibitors. To this end, we used afatinib (0.5 μM), a kinase inhibitor that irreversibly inhibits EGFR and the related ERBB2 receptor^32^, in combination with SD36 (0.5 μM), a STAT3 selective PROTAC^33^. SD36 inhibits the growth of human hematopoietic tumor cell lines and achieves effective degradation of STAT3 in tumor cells^33^. Interestingly, SD36 does not affect expression of the related STAT1 and STAT5 transcription factors (Figure 4A,B). Finally, this degrader when used in combination with afatinib, effectively induced the apoptotic death of KPeFC;*Raf*1^-/-^ tumor cells (Figure 4C-F).

**Figure 4.**
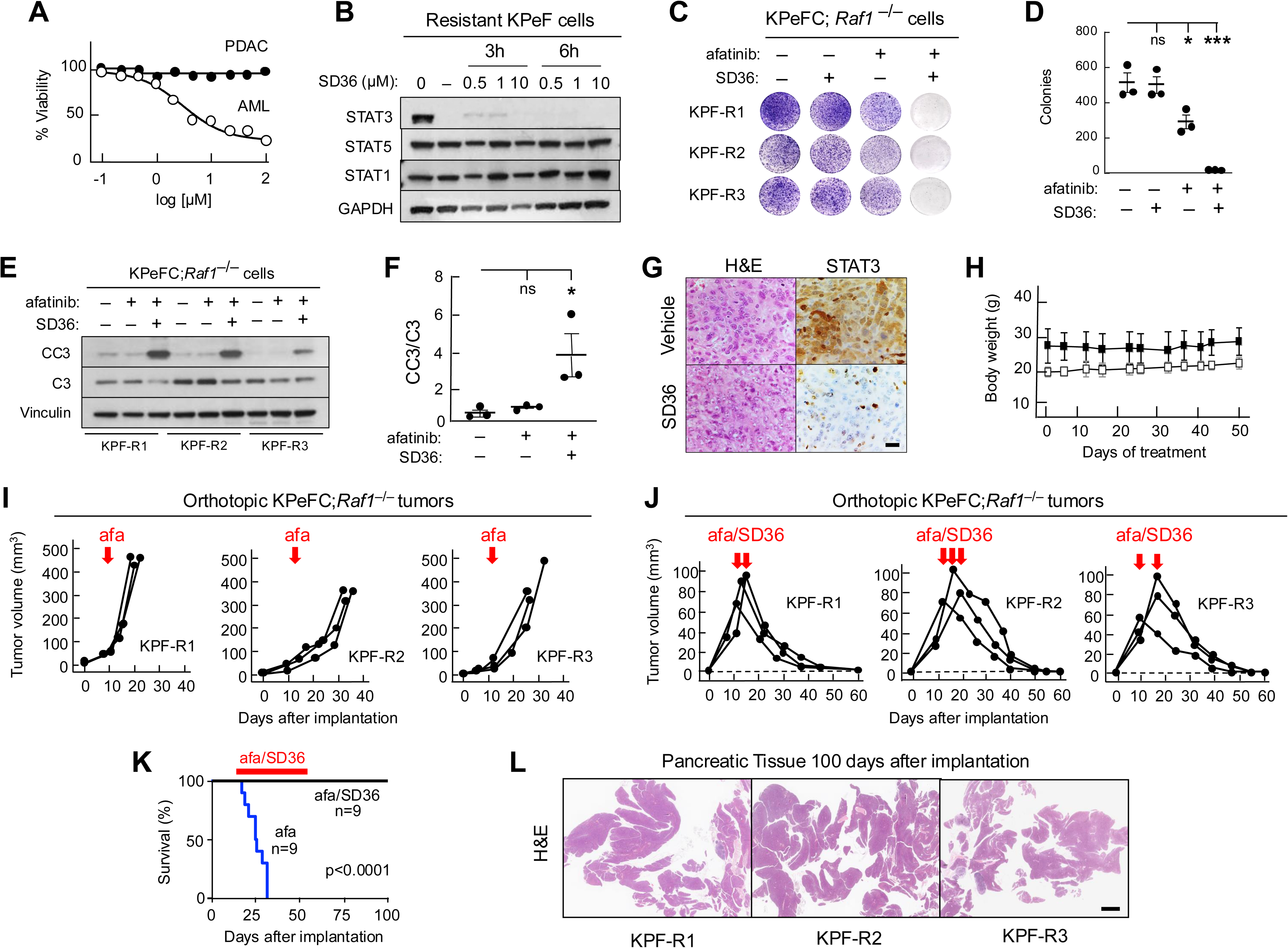
Pharmacological targeting of mouse orthotopic tumors. **A,** Effect of SD36 on the viability of representative resistant KPeF pancreatic tumor cell lines (PDAC) (solid circles) (n=3) and SU-DHL1, a human AML tumor cell line (open circles) (n=3). Viability was measured after 72h of treatment. Error bars indicate mean ± SEM. **B**, Western blot analysis of STAT3, STAT1 and STAT5 expression in whole cell extracts of a representative RAF1/EGFR resistant KPeFC;*Raf1*^L/L^;*Egfr*^L/L^ tumor cell line exposed to SD36 at the indicated concentrations and times. GAPDH was used as loading control. **C,** Colony formation assay of three representative RAF1/EGFR resistant KPeFC;*Raf1*^-/-^ tumor cell lines (KPF-R1 to KPF-R3) treated with afatinib (0.5 μM) and SD36 (0.5 μM) as indicated, for 10 days. **d**, Quantification of the number of colonies shown in **C**. The P value was obtained using multiple unpaired t tests. ns not significant, * P<0.05 and *** P<0.001. Error bars indicate mean ± SEM. **e**, Western blot analysis of CC3 and C3 in the cell lines shown in **C** and exposed to afatinib (0.5 μM) and SD36 (0.5 μM) for 24 h. Vinculin was used as loading control. **f**, Quantification of the relative expression levels of CC3 and C3 in **E**. The P value was obtained using multiple unpaired t tests. ns not significant and * P<0.05. Error bars indicate mean ± SEM. **G**, H&E and IHC analysis of STAT3 expression in sections of a RAF1/EGFR resistant KPeFC;*Raf*1^L/L^;*Egfr*^L/L^ tumor treated with either vehicle or SD36 (50 mpk, ip, qd). Scale bar represents 100 μm. Mice were treated for three days and sacrificed three hours after the end of the treatment. **H**, Body weight of tumor-bearing C57BL/6 female (n=3) (open squares) and male (n=3) (solid squares) mice treated with afatinib (20 mpk, po, qd) and SD36 (50 mpk, ip, qd) for the indicated periods of time. Error bars indicate mean ± SD. **I**, Tumor volume visualized by ultrasound of orthotopic tumors of C57BL/6 mice (n=3) implanted with three independent KPeFC;*Raf1*^-/-^ tumor cell lines (KPF-R1 to KPF-R3) treated with afatinib (20 mpk, po, qd) when the implanted tumors reached 60 to 100 mm^3^ in size (red arrows) until the tumors reached humane endpoint. **J**, Tumor volume visualized by ultrasound of orthotopic tumors of C57BL/6 mice (n=3) implanted with three independent KPeFC;*Raf1*^-/-^ tumor cell lines (KPF-R1 to KPF-R3) treated with afatinib (20 mpk, po, qd) and SD36 (50 mpk, ip, qd) when the implanted tumors reached 60 to 100 mm^3^ (red arrows) for 50 days. Dotted line represents tumor sizes undetectable by ultrasound. **K,** Kaplan-Meier survival curve of C57BL/6 mice implanted with three independent KPeFC;*Raf1*^-/-^ tumor cell lines exposed to afatinib alone (20 mpk, po, qd) (n=9) (blue line) or to a combination of afatinib (20 mpk, po, qd) and SD36 (50 mpk, ip, qd) (n=9) (solid line) for the indicated period of time (red line). Animals were kept for up to 100 days port-implantation without further treatment. The P value was obtained using Log-rank Mantel-Cox test, p<0.0001. **L**, Representative images of H&E-stained sections of pancreatic tissue of mice exposed to afatinib and SD36 and sacrificed 100 days post-implantation. No tumoral or stromal tissue could be detected. Scale bar represents 1 mm.

We nest extended these observations to *in vivo* analysis using orthotopic tumors. First, we illustrated that systemic degradation of STAT3 did not induce the toxic effects observed upon systemic ablation of S*tat*3 (Figure 4G,H). Moreover, tumor bearing mice were treated with afatinib (20 mpk, po, qd) and SD36 (50 mpk, ip, qd) once they reached 80 to 100 mm^3^ in size. Whereas mice treated with vehicle or with afatinib alone had to be sacrificed at humane endpoint due to their large tumor burden (Figure 4I and Figure S4D), those treated with the drug combination regressed rapidly until they became undetectable by ultrasound within three to four weeks (Figure 4J). Importantly, this combined treatment was well tolerated as illustrated by the absence of weight loss (Figure 4H) and by a well-preserved intestinal epithelium (Figure S4E). Importantly, at the end of the study, 100 days after implantation, all afatinib/SD36 treated mice (n=9) were alive and tumor free by ultrasound analysis (Figure 4K). Moreover, careful histopathological analysis of their pancreas at this time failed to reveal the presence of tumoral and stromal tissue, indicating that all tumors had been eliminated (Figure 4L).

### The RAS(ON) inhibitor daraxonrasib cooperates with afatinib and SD36 to induce the complete regression of orthotopic murine PDACs

Next, we interrogated whether KRAS inhibition could induce PDAC regression in the presence of RAF1 expression. *In vitro* studies revealed that the RAS(ON) inhibitor daraxonrasib effectively inhibited ERK phosphorylation as well as proliferation of sensitive and resistant tumor cells (Figure S5A-D)^15,16^. Combination of daraxonrasib (1nM) with afatinib (0.5 μM) and SD36 (0.5 μM) resulted in complete cell death, a result similar to that obtained using tumor cell lines lacking RAF1 expression (Figure S5C,D). *In vitro* combination treatments with two inhibitors, including daraxonrasib/afatinib and daraxonrasib/SD36, resulted in partial inhibition of cell proliferation whereas the combination of afatinib/SD36 was the least efficient (Figure S5C,D).

*In vivo* studies using orthotopic models derived from genetically engineered mouse (GEM) PDAC tumors led to similar results. Mice bearing tumors ranging in size from 100 to 200 mm^3^ were exposed to daraxonrasib for 30 days (20 mpk, po, qd) or until tumors reached humane endpoint. Under these conditions tumors displayed effective ERK inhibition (Figure S6A,B). Yet, as illustrated in Figure 5A, these tumors experienced partial growth inhibition that resulted in a significant increase in survival (Figure 5C). These results are reminiscent of those reported by Wasko et al using the RAS(ON) inhibitor RMC-7977 in similar orthotopic models^14^. Yet, this cohort of mice ended up succumbing to the disease after 20 to 30 days of treatment (Figure 5C). In contrast, mice treated with the same dose of daraxonrasib along with afatinib (20 mpk, po, qd) and SD36 (50 mpk, ip, qd) underwent apoptotic cell death that resulted in significant levels of tumor regression (Figure 5E,F). These tumors ended up regressing completely 20 to 30 days after initiating the combination therapy (Figure 5B,D). Moreover, these mice remained tumor free for over 200 days post-treatment, indicating that this triple therapy prevented the induction of tumor relapse (Figure 5B,C). In contrast, exposure of these tumor bearing mice to similar treatments made of dual drug combinations including daraxonrasib/afatinib, daraxonrasib/SD36 and afatinib/SD36 did not result in tumor regressions (Figure S5E). These results illustrate that concomitant targeting of these independent nodes rather than inhibiting KRAS alone, leads to the rapid disappearance of orthotopic mouse PDACs. Moreover, this triple combination also prevented the onset of tumor resistance, at least during the length of the study (Figure 5B,C). Importantly, the combined daraxonrasib/afatinib/SD36 treatment was well tolerated by the mice as determined by the lack of weight loss (Figure S6C), normal metabolic parameters, blood cell counts as well as by the integrity of several tissues including the intestinal epithelium, kidney, lung and spleen (Figure S6D-F).

**Figure 5.**
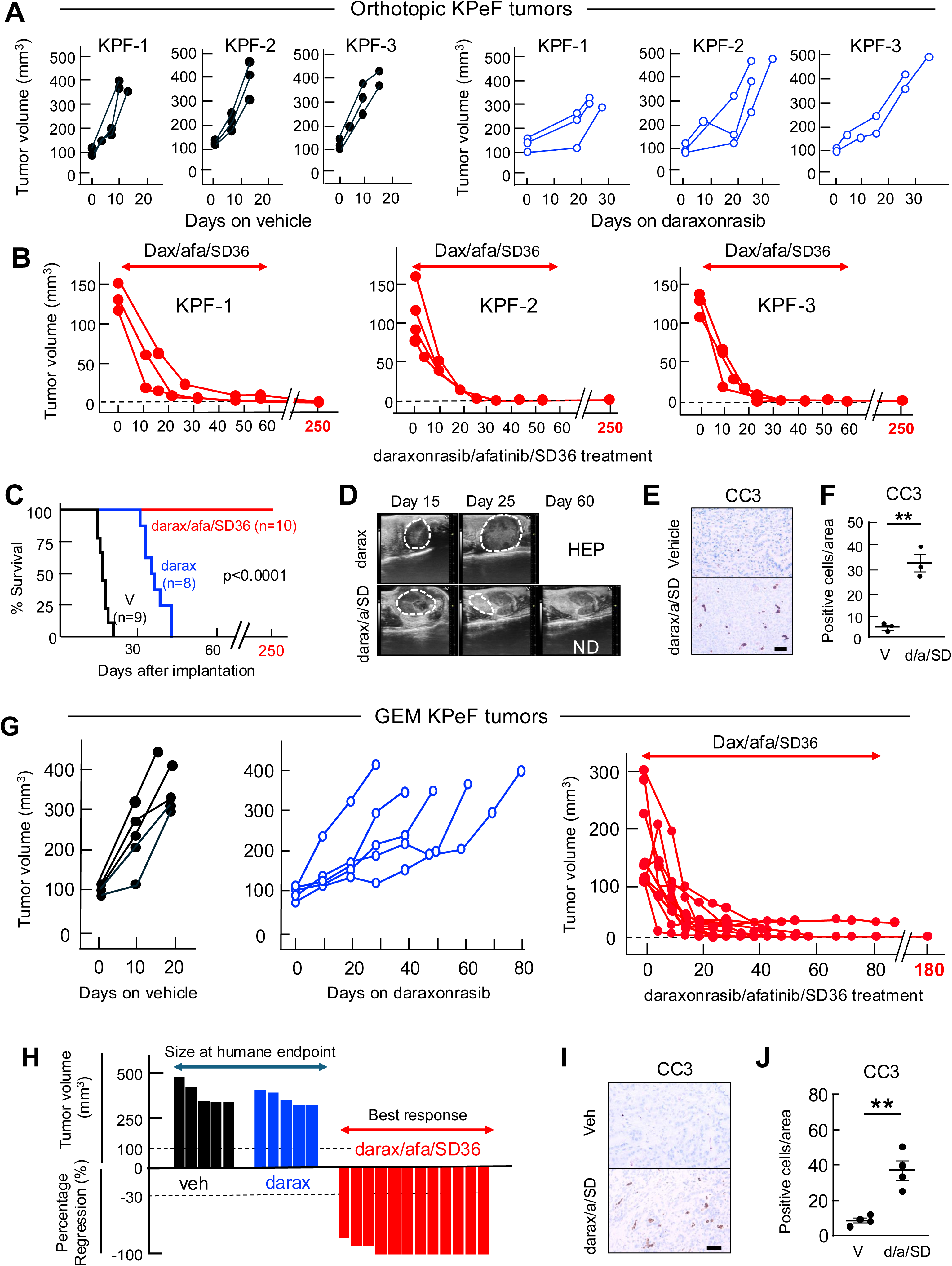
Combined inhibition of KRAS, EGFR and STAT3 in orthotopic and GEM mouse tumor models. **A**, Tumor volume visualized by ultrasound of orthotopic tumors of C57BL/6 mice implanted with three independent KPeF tumor cell lines (KPF-1 to KPF-3) treated with vehicle (solid lines) (n=9) or with the RAS(ON) inhibitor daraxonrasib (20 mpk, po, qd) (blue lines) (n=8). **B**, Tumor volume visualized by ultrasound of orthotopic tumors of C57BL/6 mice implanted with three independent KPeF tumor cell lines (KPF-1 to KPF-3) treated with a combination of daraxonrasib (20 mpk, po, qd), afatinib (20 mpk, po, qd) and SD36 (50 mpk, ip, qd) when the implanted tumors reached 120 to 200 mm^3^ in size (red arrows) for up to 60 days post-implantation (n=10). Mice were allowed to thrive for 190 additional days without further treatment. Dotted line represents tumor sizes undetectable by ultrasound. **C,** Kaplan-Meier survival curve of C57BL/6 mice implanted with three independent KPeF tumor cell lines and treated with vehicle (n=9) (V, solid line), RAS(ON) inhibitor daraxonrasib (n=8) (blue line) and a combination of daraxonrasib, afatinib and SD36 (n=10) (red line) for the indicated times. The thick lines at the top of the graph represent the periods of treatment with daraxonrasib (blue line) or with the combination of daraxonrasib, afatinib and SD36 (red line). These mice were maintained for up to 250 days post-implantation without signs of tumor relapse. The P value was obtained using Log-rank Mantel-Cox test, p<0.0001. **D,** Representative ultrasound images of the peritoneal cavity of C57BL/6 mice implanted with the KPF-1 tumor cell line and treated with daraxonrasib (darax) or with the combination of daraxonrasib, afatinib and SD36 (darax/a/SD) described above at the indicated times. Tumors are marked with dotted lines. ND, Not Detected. HEP, mouse sacrificed at humane endpoint due to large tumor burden. **E**, IHC analysis of CC3 expression in tumors of around 100 mm^3^ (n=3) exposed for four days to vehicle or to the triple combination, daraxonrasib, afatinib and SD36 and sacrificed three hours after treatment. Scale bar represents 50 μm. **F**, Quantification of the number of CC3 positive cells per area in depicted in **C**. The results represent the average of CC3 in three independent tumors. The P value was obtained using unpaired two-tailed t test. ** P<0.01. Error bars indicate mean ± SEM. **G**, Tumor volume visualized by ultrasound of GEM tumors developed by KPeF mice treated with vehicle (solid lines) (n=5), with the RAS(ON) inhibitor daraxonrasib (20 mpk, po, qd) (blue lines) (n=5) and with a combination of daraxonrasib (20 mpk, po, qd), afatinib (20 mpk, po, qd) and SD36 (50 mpk, ip, qd) (red lines). Treatment started when tumors reached at least 100 mm^3^ (0 time points). Dotted line represents tumor sizes undetectable by ultrasound. **H**, Waterfall plot of tumor response of the data shown in **G**. Tumors sizes of those tumors exposed to vehicle and those treated with daraxonrasib in monotherapy were determined ah humane endpoint. Those treated with the triple combination are represented as sizes corresponding to their best response. **I**, IHC analysis of CC3 in tumors of around 100 mm^3^ in size (n=4) exposed for ten days to vehicle or the triple combination, daraxonrasib, afatinib and SD36 and sacrificed three hours after treatment. Scale bar represents 50 μm. **J**, Quantification of the number of CC3 positive cells per area in the sections depicted in **I**. The results represent the average of CC3 in three independent tumors. The P value was obtained using unpaired two-tailed t test. ** P<0.01. Error bars indicate mean ± SEM.

Finally, the identification of the FYN kinase as a functional activator of STAT3 led us to interrogate the possibility that dasatinib, a well characterized inhibitor of the SRC family of kinases, may serve as a surrogate of the STAT3 PROTAC, SD36. Indeed, dasatinib efficiently inhibited SRC activation as determined by the inhibition of SRC^Y419^ expression both *in vitro* and *in vivo* (Figure S7A,B). In addition, dasatinib inhibited the proliferation of Resistant tumor cell lines, especially those lacking RAF1/EGFR expression (Figure S7C). Moreover, the combination of dasatinib with daraxonrasib and afatinib also induced the robust inhibition of pancreatic tumor cell proliferation *in vitro* (Figure S7D,E). Unfortunately, *in vivo* studies revealed that this triple combination was extremely toxic. As illustrated in Figure S7F,G, mice treated with this combination displayed gastrointestinal haemorrhages that led to the death of the animals within 24 hours post-treatment. Similar toxic results were obtained with related SRC family inhibitors including bosutinib, ponatinib and tirbanibulin.

### Therapeutic efficacy of combined daraxonrasib, afatinib and SD36 on GEM models of PDAC

Next, we extended the data obtained with orthotopic models to a more physiological scenario based on GEM tumor models^21^. KPeFC mice were monitored until they develop tumors ranging from 100 mm^3^ to 300 mm^3^. Mice were randomized into three cohorts and treated with either vehicle, daraxonrasib alone or with a combination of daraxonrasib, afatinib and SD36. The results obtained with the daraxonrasib arm in monotherapy resembled those obtained using orthotopic tumor models (Figure 5G and ref. 14). Of importance, mice treated with the triple combination of daraxonrasib, afatinib and SD36 displayed efficient regression of all the tumors (n=12) enrolled in the study, including nine tumors that completely disappeared within 30 to 60 days of treatment (Figure 5G,H and Figure S8). As expected, tumor regression was mediated by apoptosis (Figure 5I,J). None of these mice experienced tumor relapse at least for 80 to 180 days (Figure 5G and Figure S8).

### Inhibition of EGFR and STAT3 cooperates with RAS inhibitors in PDX tumor models

The efficacy of the triple therapy described on tumor models of human origin was evaluated using Patient-Derived Organoids (PDOs)^34,35^ and Patient-Derived Xenograft (PDX)^36^ tumor models using two different inhibitors, MRTX1133, a selective KRAS^G12D^ inhibitor^12,19^ and daraxonrasib^16^. Whole exome sequencing of the primary PDOs revealed mutations in *TP53*, *CDKN2A* and *SMAD4* tumor suppressors (Figure 6A, Figure S9A and Supplementary Table 2). PDOs were exposed for nine days to a combination of MRTX1133 (0.1 μM), afatinib (0.1 μM) and SD36 (1 μM). As illustrated in Figure 6B and Figure S9B, this treatment prevented the proliferation of tumor cells, leading to efficient cell death of all PDOs tested.

**Figure 6.**
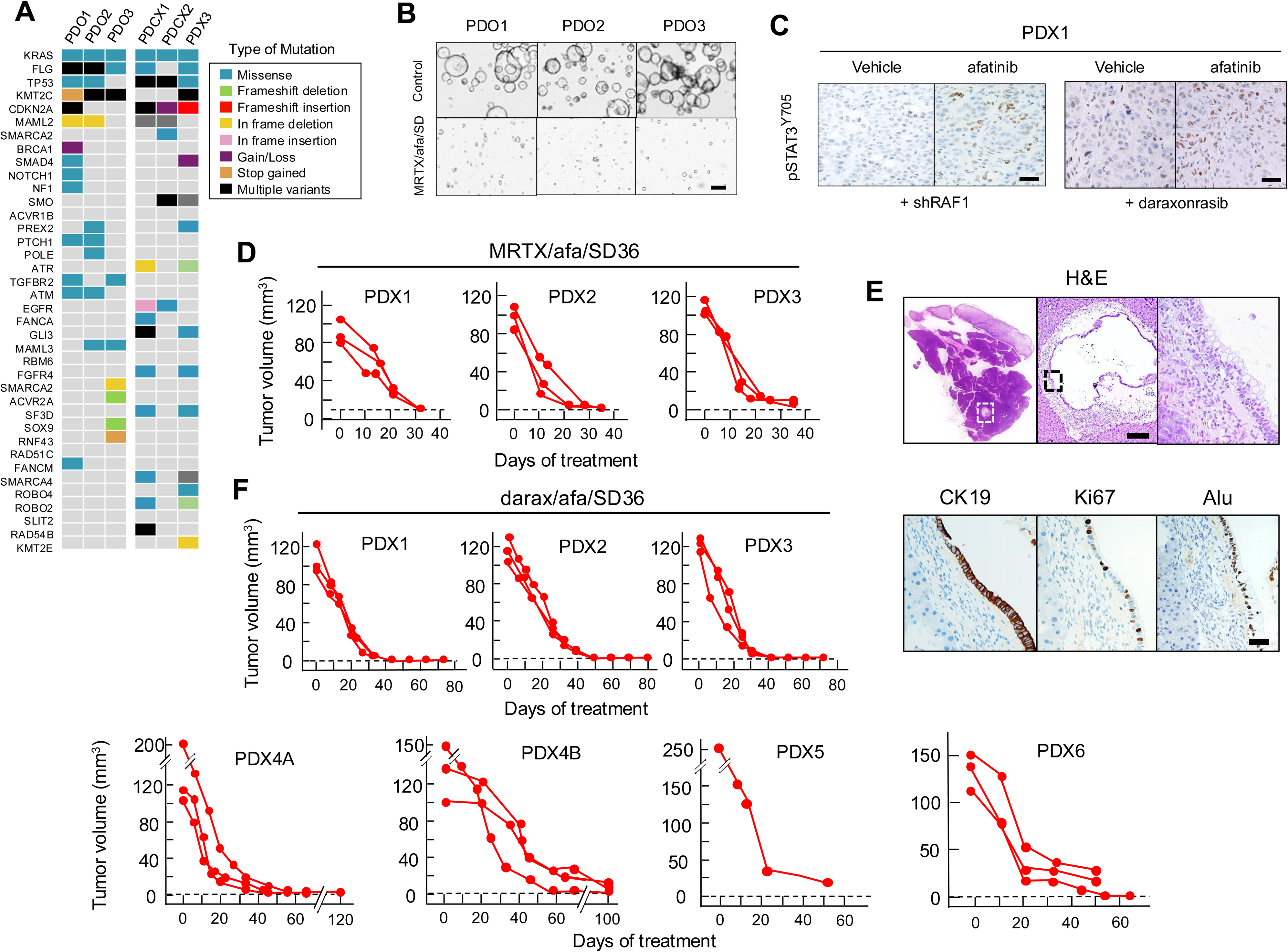
Therapeutic effect of the combined inhibition of KRAS, EGFR and STAT3 in Patient-Derived Xenografts. **A,** Oncoprint indicating the presence of mutated genes in Patient-Derived Organoides (PDO) and Patient-Derived Xenografts (PDX) analyzed by Whole Exome Sequencing. Box indicates the type of mutations. **B,** Representative images obtained with an IncuCyte® live-cell analysis system of cultures of human pancreatic organoids (PDO1 to PDO3) on day 9 of culture exposed to vehicle or to a triple combination made of the KRas^G12D^ inhibitor MRTX1133 (0.5 μM), afatinib (0.5 μM) and SD36 (1 μM). Scale bar represents 200 μm. **C,** IHC analysis of pSTAT3^Y705^ expression in the PDX1 tumor model constitutively expressing shRNAs against *Raf*1 treated for ten days with either vehicle or afatinib (20 mpk, po, qd) or treated with daraxonrasib (20 mpk, po, qd) plus either vehicle or afatinib (20 mpk, po, qd). **D**, Tumor volume of PDX1, PDX2 and PDX3 tumor models orthotopically implanted in immunodeficient mice (n=3) and visualized by ultrasound. Mice were treated for 35 days with a combination of MRTX1133 (30 mpk, ip, qd), afatinib (20 mpk, po, qd) and SD36 (50 mpk, ip, qd). Treatment was initiated when tumors reached 100 to 120 mm^3^ (0 time points). Vehicle treated tumors are represented in Figure S9C. **E,** (**Top**) Representative H&E sections of a pancreas treated with MRTX1133(Afatinib/SD36 as indicated in D (Scale bar represents 2 mm), a cyst (Scale bar represents 250 μm) and magnified section of a region of the cyst (Scale bar represents 50 μm) shown to the left. Dotted squares indicate the location of the cyst and the region magnified at the right. **(Bottom)** Representative images of CK19, Ki67 and Alu sequences of the cyst shown above. Scale bar represents 50 μm. **F**, (**Top**) Tumor volume of PDX1, PDX2 and PDX3 tumor models expanded *in vitro*, implanted in immunodeficient mice (n=3) and visualized by ultrasound. **(Bottom)** Tumor volume of PDX4A, PDX4B, PDX5 and PDX6 tumor models expanded *in vivo*, implanted in immunodeficient mice and visualized by ultrasound. Mice were treated with a combination of daraxonasib (20 mpk, po, qd), afatinib (20 mpk, po, qd) and SD36 (50 mpk, ip, qd). Treatment was initiated when tumors reached 100 to 120 mm^3^ (0 time points) and maintained for up to 120 days since the start of the treatment.

To study the effect of the triple therapy *in vivo*, we generated seven independent PDX-tumor models, including PDX1 and PDX2 whose cells had been maintained *in vitro* for an extended number of passages, PDX3 and PDX4A that consist of PDX-derived cells exclusively grown *in vitro* for expansion purposes (2-3 passages) and PDX4B, PDX5 and PDX6, three PDX models whose cells were never cultivated *in vitro* since they were directly expanded by subcutaneous implantations in immunocompromised mice. First, we validated whether these human tumor cells also underwent pSTAT3^Ty705^ activation upon *RAF1* inhibition along with afatinib exposure. As illustrated in Figure 6C, inhibition of RAF1 and EGFR led to increased expression of STAT3^Y705^ as previously observed in mouse tumors. Likewise, combined treatment of these human tumor cells with afatinib and daraxonrasib also resulted in increased expression pSTAT3^Ty705^. Thus, indicating that RAF1 and KRAS inhibition are equally effective in activating STAT3 via Tyr705 phosphorylation in human tumor cells (Figure 6C). Next, we implanted these PDX models in immunocompromised mice exposed to vehicle when they reached 100 to 120 mm^3^ in size. These mice had to be sacrificed 30 to 40 days after implantation due to their high tumor burden (Figure S9C). Histological analysis revealed high levels of cellular heterogeneity and a high percentage of proliferative cells and mitotic figures (Figure S9D). In contrast, when mice carrying similar PDX tumors were exposed to a combination of MRTX1133 (30 mpk, ip, qd), afatinib (20 mpk, po, qd) and SD36 (50 mpk, ip, qd), they underwent complete tumor regression in less than 30 days (Figure 6D). Lastly, detailed examination of the pancreas of the treated mice failed to reveal tumor or desmoplastic cells. However, six mice contained small cysts (1-10 mm^3^) lined with low proliferative human ductal-like epithelial cells without atypia (Figure 6E). Replacement of the selective KRAS^G12D^ inhibitor MRTX1133 by daraxonrasib also led to complete tumor regression in the 21 mice implanted with the seven PDX models described above (Figure 6F). Thus far, none of these animals has developed tumor resistance.

In a series of related experiments, combinations of just two of the three inhibitors/degrader failed to induce significant levels of anti-tumor activity in three independent PDX tumor models (Figure S10). As previously observed with tumors of murine origin, exposure of these human tumor cells to the triple combination also led to the appearance of apoptotic cells (Figure S11A,B). Finally, the combination therapy using either MRTX1133 or daraxonrasib, along with afatinib and SD36 was well tolerated by the immunocompromised mice as determined by body weight, histological examination of their intestinal epithelium, blood cell counts and metabolic parameters (Figure S11C-F). Thus, suggesting that mature immune T cells do not appear to be necessary for the efficient induction of tumor regression.

## DISCUSSION

Human and mouse PDAC tumors display distinct transcriptional profiles that correlate with the classical and basal subtypes^21,22,37^. Moreover, the identification of intra-tumoral heterogeneity as well as cellular plasticity it is likely to have a significant impact in predicting the efficacy of forthcoming therapeutic strategies^38^. Until recently, all therapeutic strategies against PDAC were based on the use of cytotoxic compounds. Fortunately, during the last few years, several RAS inhibitors have been developed^7-20^ including those selective against the KRAS^G12C^ and KRAS^G12D^ isoforms as well as RAS(ON) inhibitors capable of blocking additional isoforms as well as the wild type RAS proteins. These RAS(ON) inhibitors have led to significant tumor regressions and increased survival in mouse tumor models^12-20^ as well as in the clinic^,16,17^. Regrettably, the therapeutic efficacy of these inhibitors has been thwarted by the rapid appearance of tumor resistance^39^. Hence, it will be necessary to develop novel combination therapies that could induce more complete tumor regressions and limit the appearance of tumor relapses^40,41^.

We reasoned that the induction of tumor resistance could be overcome by targeting independent nodes that control KRAS signalling pathways. Indeed, we had already reported the regression of a subset of small GEM PDACs by concomitant ablation of *Raf1* and *Egfr*, two genes encoding downstream and upstream KRAS effectors, respectively^21^. As illustrated here, the tumor inhibitory response to *Raf1* and *Egfr* ablation faded away during tumor progression. Moreover, we provide experimental evidence indicating that tumor resistance involves the functional activation of the transcription factor STAT3 via phosphorylation on its regulatory residue, Tyr^705^. Indeed, resistant tumors lacking RAF1/EGFR expression undergo rapid cell death upon genetic ablation or pharmacological degradation of STAT3. Moreover, ectopic expression of a constitutively active isoform of STAT3 induced resistance to RAF1/EGFR elimination in otherwise sensitive tumor cells.

These observations led us to investigate the therapeutic effect of concomitantly ablating the three signalling nodes, R*af*1, E*gfr* and S*tat*3. As illustrated here, this strategy led to the rapid regression of pancreatic tumors. Equally important, these animals remained tumor free for over 200 days indicating that this therapeutic strategy also prevented the onset of tumor resistance over extended periods of time. Interestingly, pancreatic tumor cells maintain their oncogenic properties as long as they retain expression of one of the three signalling nodes. These observations suggest the existence of a complex signalling network that maintains the oncogenicity of pancreatic tumor cells as long as one of these independent nodes remains functionally active. Previous studies have implicated STAT3 in the resistance to MEK inhibitors. However, combination of MEK and STAT3 inhibitors only delayed tumor progression^42^. Moreover, this feedback loop appears to be mediated by the JAK kinases, suggesting that STAT3 activation is not mediated by the MAPK pathway^43^.

The absence of selective RAF1 inhibitors initially prevented us from translating those results induced by the concomitant ablation of RAF1, EGFR and STAT3 expression to a pharmacological scenario. Replacement of EGFR and STAT3 ablation by selective EGFR family inhibitor afatinib and the STAT3 degrader SD36 in mice lacking RAF1 expression faithfully reproduced the genetic results, thus validated the use of these inhibitors. Moreover, whereas systemic S*tat3* ablation was lethal, STAT3 degradation was well tolerated, thus highlighting the advantages of using reversible protein degradation versus irreversible STAT3 ablation. These results encouraged us to replace RAF1 ablation by selective RAS(ON) inhibitors. This strategy not only allowed us to complete the transition from a genetic to a pharmacological therapy but also to directly target KRAS, the initiating oncogene in PDAC.

RAS(ON) inhibitors used as monotherapy have been shown to induce partial regression of KRAS-driven orthotopic and GEM mouse tumors doubling survival of the treated mice^14-16^. These tumors, however, ended up progressing with time due to the onset of tumor resistance, ultimately leading to the death of the animals. In this study, we have confirmed these results using similar, albeit not identical tumor models (KRAS^G12V^ *vs*. KRAS^G12D^ and P53^F/F^ *vs.* P53^+/R172H^), the RAS(ON) inhibitor (RMC-6236/daraxonrasib *vs.* RMC-7977) and the dosing protocol (20 mpk, po, qd *vs.* 50 mpk, po, qod). As illustrated here, daraxonrasib monotherapy also increased survival but did not prevent the death of the animals. In contrast, our triple combination therapy based on the use of a RAS inhibitors such as daraxonrasib or MRTX1133, along with the EGFR family inhibitor afatinib and the STAT3 degrader SD36 led to complete regression of all orthotopic tumors as well as preventing the onset of tumor resistance. Indeed, treated mice remained tumor free for more than 250 days post-treatment with no signs of tumor relapse as determined by careful histological examination of mice sacrificed at the end of the experiment. Similar results were obtained with GEM tumors in which nine out of the twelve treated tumors regressed completely during the 60 days of treatment, with the remaining three displayed significant (>80%) regressions. These results, taken together, illustrate that the combination therapy of daraxonrasib/afatinib/SD36 was clearly superior to daraxonrasib monotherapy.

Genetic and biochemical evidence indicates that STAT3 activation via phosphorylation of its Tyr^705^ residue can be mediated by the FYN kinase rather than by its canonical pathway involving IL6 and the JAK kinases^26,44,45^. Unfortunately, the combination of the SRC family inhibitor dasatinib with daraxonrasib and afatinib induced gastrointestinal haemorrhages that led to the rapid death of the animals. These toxicities appeared to be mechanism based rather than drug related since we obtained similar results with other related inhibitors such as bosutinib, ponatinib and tirbanibulin. Neither of these SRC inhibitors induced significant toxicities as single agents. Further studies will be necessary to determine the mechanism(s) responsible for the observed hemorrhages.

Finally, our triple combination therapy was equally effective in inducing tumor regression in PDX tumor models. Replacement of daraxonrasib by the KRAS^G12D^ selective inhibitor MRTX1133 also led to complete regressions, indicating that the triple combination maintains its therapeutic activity with different RAS inhibitors. Moreover, these results also shed light on the mechanisms involved in tumor regression. Since PDX tumors grown in immunodeficient NU-Foxn1^nu^ mice regressed efficiently implies that the triple therapy described here exerts its antitumor effects independently of the adaptive immunity provided by the tumor microenvironment.

These observations, taken together, open the door to design novel targeted therapies for their potential application to cancer patients based on combination therapies rather than on monotherapy treatments. Indeed, the use of drug combinations is a concept that is gaining support among the RAS community^40,41^. Yet, the road to the clinic will require optimization of the combination therapy described here. For instance, a different STAT3 degrader, KT-333, is already in in clinical trials (NCT05225584) for haematological tumors. Indeed, KT-333 appears as effective as SD36 in preliminary studies. Moreover, a recently developed STAT3 PROTAC, SD-436, with improved chemical stability over SD36, may be a potential candidate for clinical trials^46^. Likewise, the role of EGFR inhibitors other than afatinib, will have to be evaluated^47^. Finally, we must keep in mind that therapeutic approaches well tolerated by mice may eventually induce unacceptable toxicities in cancer patients. Yet, despite these limitations, the results reported here should open the door to the new therapeutic options that might improve the clinical outcome of PDAC patients in a not-too-distant future.

## METHODS

### Mice

Elas-tTA/TetO-Flpo;*Kras*^+/FSFG12V^,*Trp53*^F/F^;*Raf1*^L/L^*;Egfr*^L/L^;*Rosa26*-CreERT2 mice have been previously described^21^. The conditional *Stat3*^L^ allele has also been reported^25^. Both female and male mice were used for all experiments. Immunodeficient NU-Foxn1^nu^ mice (females, 5 weeks old) were purchased from Envigo (Indianapolis, USA). All animal experiments were carried out in the animal facility of the CNIO and were approved by the Ethical Committee of the CNIO and the Institute of Health Carlos III. All strains were genotyped by Transnetyx (Tennessee, USA).

### Mouse tumor organoids

To generate mouse PDAC organoids, freshly isolated tumors were minced and digested in Advanced DMEM F12 (Gibco, USA) supplemented with HEPES 10 mM (Gibco, USA), Glutamax 1X (Gibco, USA), Collagenase P 1.5 mg/ml (Sigma, USA), Dnase I 10 μg/ml (Sigma, USA) and 0.1% BSA (Roche, Switzerland) at 37°C for 30 minutes. After resuspension in TrypLE Express (Gibco, USA), cell solutions were incubated at 37°C for 30 minutes. PDAC organoids were cultured in drops of Matrigel in Advanced DMEM F12 supplemented with HEPES 10 mM, Glutamax 1X, Rspondin1 conditioned media 10%, B27 supplement 1X (Gibco, USA), Nicotinamide 10 mM (Sigma, USA), N-acetylcysteine 1.25 mM (Sigma, USA), Noggin 0.1 μg/ml (Peprotech, USA), hEGF 0.05 μg/ml (Peprotech, USA), hFGF 0.1 μg/ml (Peprotech, USA), Gastrin I 0.01 μM (Tocris, USA) and N-2 supplement 1X (Thermo Fisher Scientific, USA). All studies were done with organoids maintained in culture for less than four passages. 20,000 cells of single-cell organoid culture were infected with 3×10^6^ PFU of Adeno-CRE viral particles in drops of Matrigel. Adeno-GFP particles were used as control. After infection, organoids were monitored in the confocal microscope for ten days after infection.

### Mouse pancreatic tumor cell lines

Mouse pancreatic explants were generated by freshly isolated PDAC tumors minced with sterile razor blades. Tumor pieces were digested with collagenase P 1.5 μg/ml in Hank’s Balanced Salt Solution (HBSS) (Gibco, USA) for 30 min at 37°C. PDAC cells were cultured in DMEM (Gibco, USA) supplemented with 10% of fetal bovine serum (FBS) (Gibco, USA) and 1X Penicillin/Streptomycin (Gibco, USA) and maintained at 37°C in humidified atmosphere with 5% CO_2_. PDAC cells were infected with Adeno-CRE viral particles (multiplicity of infection, 60) and seeded for colony formation assay 5 days after infection. Adeno-GFP particles were used as control. Cells (5,000) were seeded and allowed to form colonies for 10 days. Colonies were fixed with 0.1% glutaraldehyde (Sigma, USA) and stained with 0.5% Crystal Violet (Sigma, USA).

### Orthotopic mouse tumors

For the preparation of cell suspensions for orthotopic implantation 50,000 cells were resuspended in 50 μl of Matrigel (Corning, USA) diluted 1:1 with cold PBS (Gibco, USA). At the beginning of the surgical procedure, the analgesic Buprenorphine and the ophthalmic gel Lacryvisc (Alcon, Switzerland) were applied to mice that were anesthetized with 4% isoflurane (Braun Vetcare, Germany) in 100% oxygen at a rate of 1.5 liter/min. Bed heaters were used to avoid hypothermia due to anesthesia. After abdominal hair was removed, an incision was made in the left abdominal side, the organ of engraftment was exposed out of the peritoneal cavity and the cell suspension was injected using insulin syringes (30 gauge). The abdominal wall was sutured with absorbable Vicryl suture (Ethicon Inc., USA), and the skin was closed with wound clips (CellPoint Scientific Inc., USA). Tumor volume was assessed weekly by abdominal ultrasound. When tumors reached the appropriate size, mice were randomized and assigned into control/vehicle or treatment groups (n = 3–10 per group). Mice were either exposed to a tamoxifen-containing diet (TMX) Tekland TAM^400^/CreER (Inotiv, USA) or treated with the following inhibitors: daraxonrasib (20 mpk, po, qd), afatinib (20 mpk, po, qd) and SD36 (50 mpk, ip, qd) either alone or in the indicated combinations.

### Patient-derived organoids

Patient-Derived Organoids (PDO) were generated from tumors collected from patients selected for surgery as first-line treatment. Samples were received from the Hospital Clínico Universitario Virgen de la Arrixaca (Murcia, Spain). Specific informed consent for patient-derived organoids (PDOs) generation was obtained from all patients. Samples and data from patients included in this study were provided by the Biobank IMIB (National Registry of Biobanks B. 0000859) (PT23/00026), integrated in the Platform ISCIII Biomodels and Biobanks and were processed following standard operating procedures with the appropriate approval of the Ethics and Scientific Committees (CEIC HCUVA-2013/01).

To generate PDOs, tumor samples were chopped and incubated in advanced DMEM/F12 (Gibco, USA) complemented with HEPES 10 mM (Gibco, USA), Glutamax 1X (Gibco, USA), Primocin 100 μg/ml (Thermo Fisher Scientific, USA), BSA 0.1% (Roche, Switzerland), Collagenase XI 2.5 mg/ml (Sigma, USA), DNAse I 10 μg/ml (Sigma, USA) and Y-27632 10.5 μM (Sigma, USA) in in rotation at 37°C for two rounds of 30 minutes, as previously described^34^. Tumoral organoids were seeded in drops of Matrigel (Corning, USA) in advanced DMEM/F12, HEPES 10 mM, Glutamax 1X, A83-01 500 nM (Tocris, UK), human Epidermal Growth Factor (hEGF) 50 ng/ml (Peprotech, USA), Noggin 100 ng/ml (Peprotech, USA), human Fibroblast Growth Factor (hFGF10) 100 ng/ml (Peprotech, USA), Gastrin I 0.01 μM (Tocris, UK), N-acetylcysteine 1.25 mM (Sigma, USA), Nicotinamide 10 mM (Sigma, USA), B27 supplement 1X (Gibco, USA), Rspondin1 conditioned media 10%, and Wnt3A conditioned media 50%. PDOs were maintained at 37°C with 5% CO_2_.

### Incucyte assays

PDOs were recovered from Matrigel and prepared as single cell suspension, after incubation with trypLE Express (Gibco, USA) in RT for 30 mins. 250-500 viable cells were plated in 4 µl of Matrigel with the Matribot® Bioprinter (Corning, USA). Organoids were seeded in 96-well plates and maintained in Human Complete Feeding Medium with MRTX1133 0.1 μM (MedchemExpress, USA), Afatinib 0.1 μM (Amatek, USA) and SD36 1 μM (Domainex, UK) were added 72 hours after seeding. All conditions were tested in octuplets. PDOs viability was followed for 9 days with the Sartorius Incucyte® S×5 Live-Cell Analysis System (Sartorius, Germany). Incucyte® Organoid Analysis Software Module (v2023A) was applied to recognize PDOs and draw the viability curves by calculating the organoid individual area normalized to day 0 of treatment.

### Patient-Derived Xenografts

We have used three types of PDX tumor models including PDX1 and PDX2 (PDX models derived from long-term passaged in culture^21^, PDX3^48^ and PDX4A (PDX models exclusively grown in culture for 2-3 passages for expanding purposes) and PDX4B (a model expanded *in vivo* via subcutaneous implantations). In all cases, tumor models were generated from surgically removed samples as described above. Samples are included in registered projects that follow standard operating procedures with the appropriate approval of the Ethics and Scientific Committees (numbers CEIC HCUVA-2013/01, 2011-A01439-32 and CEI 60-1057-A068). For the generation of the PDX1 and PDX2 as well as PDX3 and PDX4A tumor models, cell suspensions (100,000 cells) were orthotopically implanted in the pancreas of immunodeficient mice as described above for the corresponding mouse tumors. In the case of the PDX4B, PDX5 and PDX6 tumor models, human tumor pieces were directly implanted subcutaneously in the flank of immunodeficient mice until tumors reached a size of 100 mm^3^. Tumor volume was assessed weekly by abdominal ultrasound (see below). When tumors reached the appropriate size, mice were randomized and assigned into control/vehicle or treatment groups (n = 3–10 per group). Mice were treated with the following inhibitors: MRTX1133 (30 mpk, ip, qd), daraxonrasib (20 mpk, po, qd), afatinib (20 mpk, po, qd) and SD36 (50 mpk, ip, qd) either alone or in the indicated combinations.

### Inhibitors

Daraxonrasib (RMC-6236), MRTX1133, dasatinib, ruxolitinib, itacitinib, tofacitinib, baricitinib, bosutinib, ponatinib, tirbanibulin and galunisertib were purchased from MedChemExpress (NJ, USA). Afatininb was obtained from Amatek (PA, USA). SD36 was synthesized by Domainex (Cambridge, UK). For *in vitro* experiments these inhibitors were diluted in DMSO to final concentration of 10 mM. For *in vivo* experiment afatinib was dissolved in 0.5% methylcellulose and 0.2% Tween-80 in water, SD36 in 10% PEG in PBS and daraxonrasib, MRTX1133 and dasatinib in 2-10% DMSO, 40% PEG300, 5% Tween-80 in saline, according to the guidelines of the manufacturer. *In vivo* experiments with bosutinib (100 mpk), ponatinib (30 mpk) and tirbanibulin (25 mpk) were performed in 10% DMSO, 40% PEG300, 5% Tween-80 in saline as vehicle solution.

### Tumor monitorization

Tumor monitorization was carried out by bi-weekly ultrasounds in the Molecular Imaging Core Unit of CNIO. Briefly, mice were anesthetized with 4% isoflurane (Braun Vetcare, Germany) in 100% oxygen at a rate of 1.5 liter/min. Bed heaters were used to avoid hypothermia due to the anesthesia and abdominal hair was locally removed. Tumors were measured with a High-Resolution system Vevo 770 (Visualsonics, Canada) in an ultrasound transducer of 40 MHz (Visualsonics RMV704, Canada). Tumor sizes were calculated as (Length*Width*Height)/2.

### Blood analysis

Blood was collected from the heart of euthanized mice and transferred to an EDTA-containing tube for hemogram analysis using the LaserCell blood counter. In parallel, blood also collected from the heart was used for analysis of the metabolic profile with a diagnostic rotor on a VetScan VS2 analyzer.

### Immunohistochemistry

Specimens were fixed in 10% formalin (Sigma) and embedded in paraffin. For histopathological analysis, specimens were serially sectioned (3 μm thick) and every ten sections stained with hematoxylin and eosin (H&E). Immunohistochemistry staining was performed by the Histopathology Unit of CNIO. Antibodies used for mouse tissue include STAT3a (D1A5) (Cell Signaling Technology 8768), EGFR (D1P9C) (Cell Signaling Technology 71655), RAF1 (EM1411E) (Monoclonal Antibodies Core Unit, CNIO), phospho-STAT3 (Y705) (Cell Signaling Technology 9145), phospho-p44/42 MAPK (Cell Signaling Technology 9101), Cleaved Caspase 3 (Cell Signaling Technology 9661) and phospho-SRC (Y419) (Abcam ab4816).

For human tissue, the antibodies phospho-STAT3 (Y705) (Cell Signaling Technology 9145), phospho-EGFR (Cell Signaling Technology 2234), Cleaved Caspase 3 (Cell Signaling Technology 9661), Cytokeratin 19 (Agilent IR615) and Ki67 (Agilent IR626) were used. For the antibodies against pSTAT3 and CC3 different dilutions and incubation times were used (1:300, 48 min for mouse tissue and 1:200, 24 min for human tissue). For the detection of Alu sequencies the nuclear marker Nucleoli (Abcam ab190710) was used. Briefly, for *in situ* hybridization, sections were deparaffinized and re-hydrated. Next, antigen retrieval was first performed with the selected buffer (ULTRA CC2, 5424542001, Roche) and discovery protease III (760-2020, Roche). Slides were then incubated with the probe AluII (05272041001 Roche) labeled with DNP, followed by stringency washes and incubation with Rabbit anti-DNP (Sigma) conjugated with horseradish peroxidase. Immunohistochemical reaction was developed using DAB+, as a chromogen (ChromoMap DAB, Roche). Finally, nuclei were counterstained with Carazzi’s hematoxylin, all the slides were dehydrated, cleared and mounted with a permanent mounting medium for microscopic evaluation. Stained slides were scanned using the <u>Mirax</u>scanner (Zeiss, Germany). Images were analyzed by the Zen2 software and photos were exported using Zen2 software (Zeiss, Germany).

### Colony formation assays

Infected cells (5,000) were seeded and allowed to form colonies for 10 days. Colonies were fixed with 0.1% glutaraldehyde (Sigma, USA) and stained with 0.5% Crystal Violet (Sigma, USA). All colony formation assays were performed three times and all plates were seeded in duplicates.

### shRNAs

Cells derived from mouse PDAC explants were infected with lentiviral supernatants expressing shRNAs against *Egfr* (TRCN0000055218, TRCN0000055220), *Raf1* (TRCN0000012628, TRCN0000055140), *Stat*3 (TRCN0000071453, TRCN0000301946), *Il6ra* (TRCN0000068293, TRCN0000305257), *Jak*1 (TRCN0000321912, TRCN0000023290), *Jak*2 (TRCN0000278125, TRCN0000278192), *Fyn* (TRCN0000361149, TRCN0000361212), *Stat5* (TRCN0000231549, TRCN0000231550), T*gftr3* (TRCN0000348460, TRCN0000334910), S*1pr1* (TRCN0000173790, TRCN0000175553) and *Rel*A (TRCN0000235832, TRCN0000235834). PDX-derived cells were infected with lentiviral supernatant expressing shRNAs against *Raf*1 (TRCN0000195646, TRCN0000197115). shRNAs were cloned in plasmids carrying a cassette of resistance to Puromycin or in plasmids that confer resistance to Blasticidine. Non-targeted shRNA vectors were used as negative controls.

Nucleotide sequences of all the shRNAs described above are indicated below:

TRCN0000055218: GCTGGATGATAGATGCTGATA

TRCN0000055220: CCAAGCCAAATGGCATATTTA

TRCN0000012628: GCTTTGGTACTACAGAACTTT

TRCN0000055140: CAAGCAATACTATCCGGGTTT

TRCN0000071453: CCTAACTTTGTGGTTCCAGAT

TRCN0000301946: GCAGGTATCTTGAGAAGCCAA

TRCN0000068293: CGAAGCGTTTCACAGCTTAAA

TRCN0000305257: TGGGTCTGACAATACCGTAAA

TRCN0000321912: GCCCTGAGTTACTTGGAAGAT

TRCN0000023290: GCCCTGAGTTACTTGGAAGAT

TRCN0000278125: CGTGGAATTTATGCGAATGAT

TRCN0000278192: CGGCCCAATATCAATGGATTT

TRCN0000361149: TCTTCACCTGATTCAACTAAA

TRCN0000361212: CACTGTTTGTGGCGCTTTATG

TRCN000012549: GCCAAGTATTACACTCCTGTA

TRCN000012550: CGCCAGATGCAAGTGTTGTAT

TRCN0000348460: GTGGTTTACTATAACTCTATT

TRCN0000334910: GCAGAGAATGAGCATGTATAT

TRCN0000173790: CATGGAATTTAGCCGCAGCAA

TRCN0000175553: CGTCTGGAAACGTCAATTCTT

TRCN0000235832: AGGCCATATAGCCTTACTATC

TRCN0000235834: GCTCAAGATCTGCCGAGTAAA

TRCN0000195646: CCAACACTCTCTACCGAAGAT

TRCN0000197115: GCTCAGGGAATGGACTATTTG

### Flow cytometry

Cells were harvested after trypsinization and washed with PBS. To assess apoptosis, cells were stained with Annexin V-APC (BioLegend, USA) and DAPI (Thermo Fisher Scientific, USA). A total of 10⁶ cells were resuspended in 100 µL of binding buffer and incubated with Annexin V and DAPI for 20 minutes at room temperature in the dark. After staining, samples were analyzed by flow cytometry using an Attune Nxt Flow Cytometer (Thermo Fisher Scientific, USA) and data was processed with FlowJo (v 10). Single cells were obtained by gating in the FSC-H/FSC-A plot and debris was removed in the FSC-A/SSC-A plot. Annexin V-positive and DAPI-negative cells were considered early apoptotic. Double positive cells were considered late apoptotic, double negative cells were considered alive, and Annexin V-negative DAPI-positive cells, necrotic.

### MTT viability assay

Cells were seeded in 96-well plates at a density of 5,000 cells/well and incubated in their respective culture medium. Treatment was started 24 hours later for 72 hours, and viability rate was determined by the 3-(4,5-dimethylthiazol-2-yl)-2,5-diphenyltetrazolium bromide (MTT) assay (Roche, Switzerland). Absorbance was measured with a microplate reader at 590 nm (EnVision 2104 Multilabel Reader, Perkin Elmer, USA). All samples were assayed in triplicates and results represent the average of viability normalized to the vehicle control. All experiments were performed at least three times.

### Reverse Transcription quantitative PCR (RT-qPCR)

Total RNA was extracted from cell pellets using the RNeasy Mini Kit (Qiagen, USA), according to manufactureŕs protocol, followed by quantification. First, 1 μg of total RNA samples was converted cDNA using the SuperScript II Reverse Transcription Kit (Thermo Fisher Scientific, USA). Next, RT-qPCR reaction was performed using the Power SYBR Green Master Mix (Applied Biosystems, USA). Results were analyzed using the 2^-ΔΔCt^ values of the mean threshold cycle (Ct) value obtained from three replicates. Primers used include *Il6ra* (Fw: AGCGACACTGGGGACTATTTA, Rv: ACAGCCTTCGTGGTTGGAG)*, Jak1* (Fw: ACGCTCCGAACCGAATCATC, Rv: GTGCCAGTTGGTAAAGTAGAACC)*, Jak2* (Fw: CTTGTGGTATTACGCCTGTGT, Rv: TGCCTGGTTGACTCGTCTATG)*, RelA* (Fw: AGGCTTCTGGGCCTTATGTG, Rv: TGCTTCTCTCGCCAGGAATAC), *Tgfbr3* (Fw: CATCTGAACCCCATTGCCTCC, Rv: CCTCCGAAACCAGGAAGAGTC)*, S1pr1* (Fw: CGCAGTTCTGAGAAGTCTCTGG, Rv: GGATGTCACAGGTCTTCGCCTT)*, Birc5* (Fw: CTACCGAGAACGAGCCTGATT, Rv: AGCCTTCCAATTCCTTAAAGCAG)*, Bcl-2* (Fw: ATGCCTTTGTGGAACTATATGGC, Rv: GGTATGCACCCAGAGTGATGC)*, Bcl-XL* (Fw: GAAAGCGTAGACAAGGAGATG, Rv: ACAAGGGGCGTGGTTCTTAC)*, Mcl-1* (Fw: CAAAGATGGCGTAACAAACTGG, Rv: CCGTTTCGTCCTTACAAGAACA), and *Gapdh.* (Fw: AGCGACACTGGGGACTATTTA, Rv: ACAGCCTTCGTGGTTGGAG) as housekeeping control.

### Western blot analysis

Protein extraction was performed on ice with NP-40 lysis buffer supplied with protease and phosphatase inhibitors (cOmplete Mini, Roche; Phosphatase Inhibitor Cocktail 2 and 3, Sigma). To quantify the concentration of protein lysates, the Bradford method (Bio-Rad) was used. 25 μg of cell extracts were loaded and wet transfer was performed using nitrocellulose transfer membrane and transfer buffer (Tris-Glycine 1X, Lonza; MeOH 20%, Panreac), for 70 minutes at a constant current of 0.40 A. Membranes were blocked by incubation with 5% non-fat milk in TBST (1X Tris-Buffered Saline (TBS) solution, 0.1% Tween-20, Sigma). Incubation with primary antibodies diluted in blocking solution was performed overnight on a rotating platform at 4°C. Next, membranes were washed thrice with 1X TBST buffer for 10 minutes on a shaking platform at RT, following incubation with HRP secondary antibodies. Protein visualization was carried out with enhanced chemiluminescent system (ECL Plus, Amersham Biosciences). Membranes were blotted with antibodies raised against EGFR (1:500, Abcam ab52894), RAF1 (1:500, BD Biosciences 610151), STAT3 (1:500, Cell Signaling Technology 9139), phospho-STAT3 (Tyr705) (1:500, Cell Signaling Technology 9131), STAT5 (1:1000, Cell Signaling Technology 94205), STAT1 (1:1000, Cell Signaling Technology 9172), phospho-p44/42 MAPK (1:1000, Cell Signaling Technology 4370), p44/42 MAPK (1:1000, Cell Signaling Technology 9107), Cleaved Caspase 3 (1:250, Cell Signaling Technology 9661), Caspase 3 (1:500, Cell Signaling Technology 9662), FYN (1:1000, Atlas Antibodies HPA023887), phospho-SRC (Y419) (1:500, Abcam ab4816), SRC (1:500, Cell Signaling Technology 2108) and GAPDH (1:5000, Sigma G8795), VINCULIN (1:1000, Sigma-Aldrich V9131), and ACTIN (1:1000, Sigma-Aldrich A5441) as loading controls. Anti-mouse HRP-linked (1:2000, Agilent Dako P0447) and anti-rabbit HRP-linked (1:2000, Agilent Dako P0448) were used as secondary antibodies. For the quantification of protein expression, the intensity of the bands was quantified with ImageJ2.

### RNA sequencing

Total RNA was extracted from mouse samples using the RNeasy Mini Kit (Qiagen, USA), according to manufactureŕs protocol, followed by assessment of the quality using the RNA Integrity factor (RIN). Library preparation and RNA Sequencing were performed in the Genomics Unit of CNIO. First, 1 μg of total RNA samples was converted into sequencing libraries with the NEBNext Ultra II Directional RNA Library Prep Kit (New England Biolabs E7760, USA), as recommended by the manufacturer. Briefly, polyA+ fraction was purified and randomly fragmented, converted to double stranded cDNA and processed through subsequent enzymatic treatments of end-repair, dA-tailing and ligation to adapters. Adapter-ligated library was completed by PCR with Illumina single-end primers. Next, purified cDNA libraries were applied to an Illumina flow cell for cluster generation and sequenced on an Illumina instrument (Illumina NextSeq 500, USA) following manufacturer’s protocols.

### Differential expression and GSEA analysis

Fastq files were processed using the Cluster_rnaseq pipeline developed by the Bioinformatic Unit of CNIO (https://github.com/cnio-bu/cluster_rnaseq). Briefly, FastQC (v 0.12.1) and MultiQC (v 1.18) were used for quality control. STAR (v 2.7.11a) and featureCounts (v 2.0.6) were used as aligner to the mouse reference genome GRCm39 and quantifier, respectively, and differential expression analysis was performed with DESeq2 (v 1.30.1). GSEAPreranked (v 4.3.2) was performed for functional enrichment of gene sets among the different groups of cells. As gene sets, the collection of Hallmark (MH) and the Reactome subset of the Canonical Pathways (CP) were selected from the Mouse Molecular Signatures Database (MsigDB) collections.

### Whole Exome Sequencing (WES)

Genomic DNA was obtained from PDO, PDCX and PDX cultures followed by quantification. Sample preparation and sequencing were performed by Novogene Ltd (UK). Briefly, after assessment of quality, the SureSelect XT_V6 Kit (Agilent Technologies, USA) was used for library preparation. Samples were applied to a NovaSeq 6000 instrument (Illumina, USA) for pair-end (PE) sequencing of 150 bp fragments.

### Variant calling and annotation

Fastq files were processed using the Varca pipeline developed by the Bioinformatic Unit of CNIO (https://github.com/cnio-bu/varca). Briefly, FastQC (v 0.12.1) and MultiQC (v 1.19) were used for quality control. Sequences were aligned to the hGRC38 reference genome with BWA-MEM (v 0.7.17). Refinement of alignment, variant calling and annotation steps were performed by Picard (v 3.0.0), GATK (v 4.0.0), MuTect2 and VEP (v 104).

### Statistical analysis

Data are represented as mean ± SEM or ± SD. Significance was calculated with paired or unpaired t test, as indicated in the figure legends, using GraphPad Prism. Log-rank Mantel-Cox test was used to compare survival curves using GraphPad Prism. Significant differences between experimental groups were indicated as following: * P< 0.05, ** P< 0.01 or *** P< 0.001.

### Data availability

Raw RNA-seq and WES data have been uploaded to the National Center for Biotechnology Information’s Gene Expression Omnibus (GEO) and Sequence Read Archive (SRA) Database repositories, with identifiers GSE271518 and PRJNA1130922 respectively. This paper does not report original code.

## ACKNOWLEDGEMENTS

We thank R. Villar for excellent technical assistance; I. Aragón, V. Viñas and I. Blanco (Animal Facility); Francisca Mulero, G. Visdomine and G. Medrano (Molecular Imaging Unit), P. Gonzalez (Histopathology Unit) and O. Dominguez (Genomic Unit) of CNIO for their excellent technical support. We also thank Jesús de la Peña and Eduardo Ortiz from the Servicio de Anatomía Patológica of the *Hospital Clínico Universitario Virgen de la Arrixaca* (HCUVA) (Murcia) and the Biobank of the *Instituto Murciano de Investigación Biosanitaria* (IMIB) (PT23/00026; Platform ISCIII Biomodels and Biobanks) for providing clinical samples. This work was supported by grants from the CRIS Cancer Foundation, the European Research Council (ERC-AG/695566-THERACAN), the *Agencia Estatal de Investigación* co-funded with the European Regional Development Fund (ERDF-EU ERDF “A way of making Europe”) (PID2021-124106OB-I00; MCIU/AEI/10.13039/501100011033) and the European Union “NextGenerationEU”/PRTR” (PLEC2022-009255; MCIU/AEI/10.13039/501100011033) to M.B. M.B. and C.G. are recipients of a CIBERONC Fund (CB21/12/00121). Additional funding included grants from the *Instituto de Salud Carlos III* co-funded by ERDF “A way of making Europe” (PI19/00514), the “*Carmen Delgado/Miguel Pérez Mateo Grants/Asociación Cancer de Páncreas*” & “*Asociación Española de Pancreatología*” to C.G., and the Italian Cancer Research Association (AIRC, IG16930) and the Italian Ministry of University and Research (MIUR PRIN 2017) to V.P. M.B. is the recipient of an Endowed Chair from the AXA Research Fund. V.L. was supported by an INPhINIT fellowship from “La Caixa” Foundation (LCF/BQ/DI18/11660011) and a contract from the CRIS Cancer Foundation. S.B. is supported by a PhD scholarship from the Foundation of Science and Technology of Portugal (2021.05875.BD). M.K. was supported by an Erasmus+ scholarship for traineeship. L.M.-C. was supported by an FPU fellowship from the *Ministerio de Educación*. J.C.L-G. is supported by a postdoctoral fellowship from *Amigos del CNIO*, E.Z.-D. is supported by a FPI fellowship (PRE2022-102952) from the Spanish Ministry of Sciences and Innovation (PID2021-124106OB-I00). P.S. is partially supported by a fellowship from the China Scholarship Council.

## AUTHOR CONTRIBUTIONS

C.G. and M.B. conceptualized the study and designed research; V.L. conducted most of the experiments; S.B., B.R-P., M.K., C.G.-L., P.S., J.C.L.-G. and E.Z.-D. contributed to *in vitro* studies; S.J.-P., R.B., S.B., A.L-G. and L.M.-C. helped with *in vivo* studies; V.L. and R.A. analyzed transcriptomic and exomic data; M.S.R. performed molecular biology studies; E.C. performed histopathology analyses, B.S.Jr. and N.D. provided PDX tumors; V.P. the *Stat3*^lox^ allele and F.S.-B. fresh human PDACs; M.M., and M.D. provided critical input;, V.L, C.G. and M.B. interpreted the data and prepared the manuscript.

## SUPPLEMENTARY INFORMATION

Supplementary Table 1. Excel file containing differentially expressed genes in Sensitive and Resistant cells.

Supplementary Table 2. Excel file containing selected variants of PDO and PDX samples.

## SUPPLEMENTAL FIGURE LEGENDS

**Figure S1.**
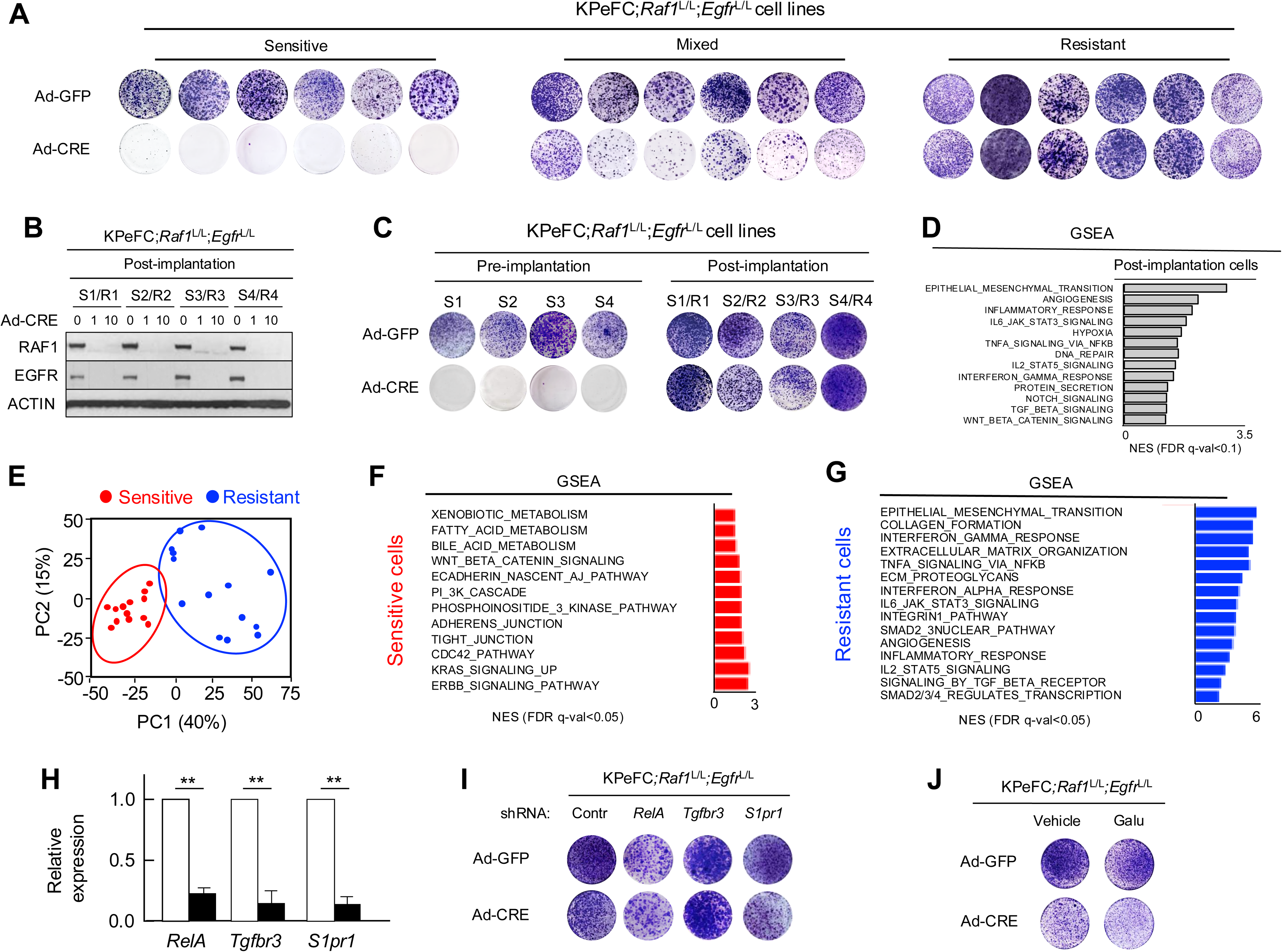
Distinct molecular profiles of Sensitive and Resistant tumor cells. **A,** Colony formation assay of representative KPeFC;*Raf1*^L/L^;*Egfr*^L/L^ tumor cell lines classified as Sensitive, Mixed or Resistant based on their response to *Raf*1 and *Egfr* after infection with Adeno-GFP and Adeno-CRE viral particles. **B**, Western blot analysis of Sensitive cells converted into Resistant cells (S1/R1 to S4/R4) to illustrate that retained the absence of RAF1 and EGFR expression. **C,** Colony formation assay of representative KPeFC;*Raf1*^L/L^;*Egfr*^L/L^ sensitive cell lines (S1 to S4) that acquired a resistant phenotype upon passage as orthotopic tumors in immunocompetent C57BL/6 mice (S1/R1 to S4/R4). **D,** GSEA analysis of post-implantation versus pre-implantation tumor cell lines. Pathways enriched in converted RAF1/EGFR Resistant tumor cells are indicated by grey bars. The Normalized Enrichment Score (NES) is represented. **E**, Principal Component Analysis (PCA) of transcriptomic data indicating the distribution of tumor cell lines Sensitive (n=13) and Resistant (n=14) to RAF1/EGFR ablation. **F,** GSEA analysis of Sensitive tumor cell lines. The Normalized Enrichment Score (NES) is represented. **G,** GSEA analysis of Resistant tumor cell lines. The Normalized Enrichment Score (NES) is represented. **H**, Expression levels of *RelA*, *Tgfbr3* and *S1pr1* genes as determined by RT-qPCR analysis in RAF1/EGFR resistant KPeFC;*Raf1*^L/L^;*Egfr*^L/L^ tumor cell lines expressing either control shRNAs (open bars) or shRNAs against the respective genes (solid bars). The P value was obtained using paired two-tailed t test. ** P<0.01. Error bars indicate mean ± SEM. **I**, Colony formation assay of a representative RAF1/EGFR resistant KPeFC;*Raf1*^L/L^;*Egfr*^L/L^ tumor cell line expressing the indicated shRNAs and infected with Adeno-GFP or Adeno-CRE viral particles. **J**, Colony formation assay of a representative RAF1/EGFR resistant KPeFC;*Raf1*^L/L^;*Egfr*^L/L^ tumor cell line treated with either vehicle or with 5 μM Galunisertib (Galu) for 10 days followed by infection with Adeno-GFP or Adeno-CRE viral particles.

**Figure S2.**
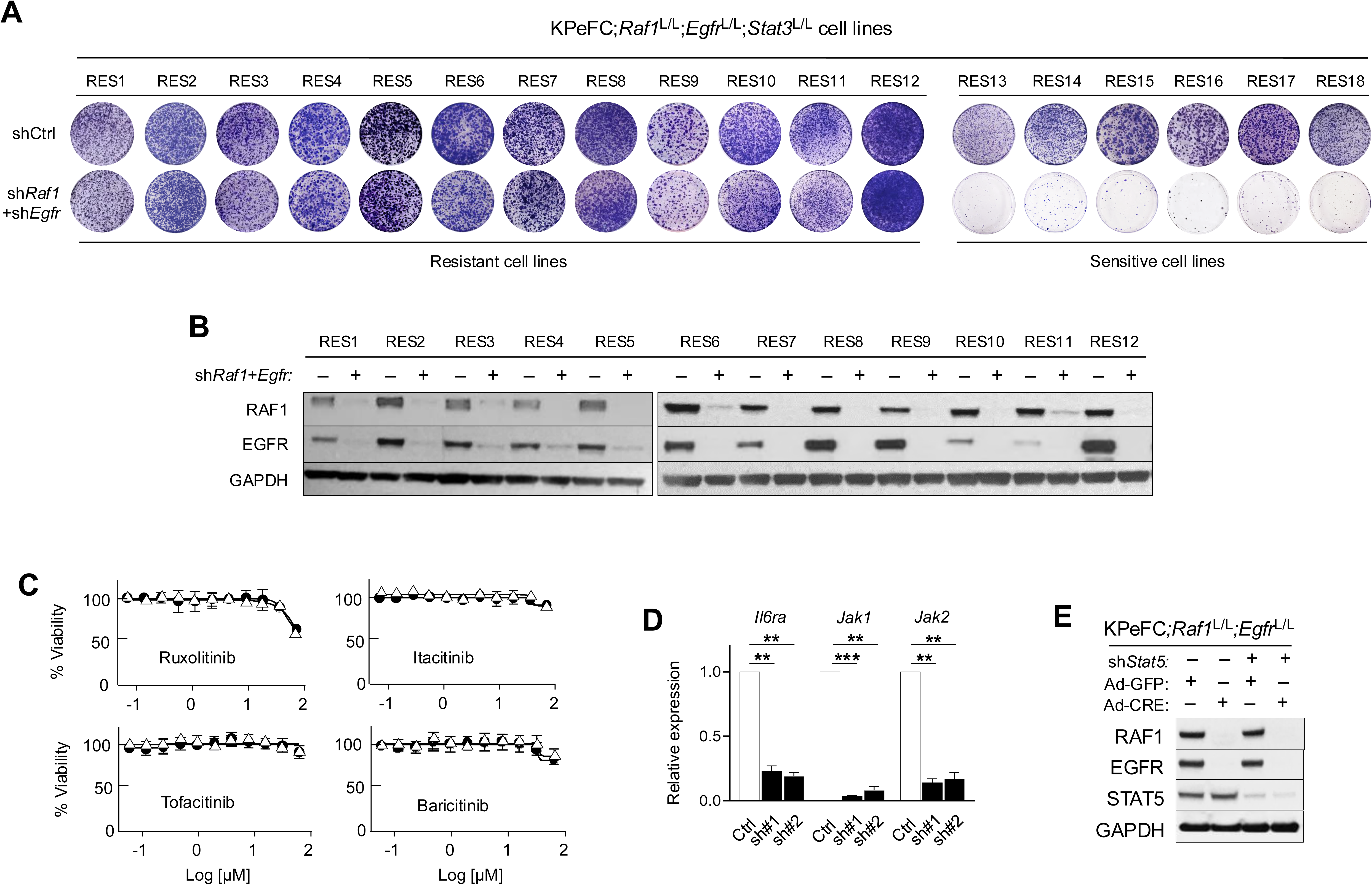
Jak1/2 inhibitors do not affect viability of PDAC tumor cell lines. **A**, Identification of KPeFC;*Raf*1;*Egfr*^L/L^;*Stat3*^L/L^ tumor cell lines Resistant (RES1 to RES12) or Sensitive (RES13 to RES18) to RAF1/EGFR expression based on their ability to proliferate in the presence of shRNAs against *Raf1* and *Egfr*. Nontargeted shRNAs were used as negative control (shCrtl). **B**, Western blot analysis of RAF1 and EGFR expression in cell extracts of representative RAF1/EGFR resistant (RES1 to RES12) KPeFC;*Egfr*^L/L^;*Raf1*^L/L^;*Stat3*^L/L^ cell lines exposed to shRNAs against *Raf1* and *Egfr*. GAPDH was used as loading control. **C,** Viability of three representative RAF1/EGFR resistant KPeFC;*Raf1*^L/L^;*Egfr*^L/L^ (solid circles) and KPeFC;*Raf1*^-/-^;*Egfr*^-/-^ (open triangles) tumor cell lines exposed to the indicated JAK1/2 inhibitors for 72 hours. Error bars represent mean ± SEM. **D**, Expression levels of I*l6ra*, J*ak1* and J*ak2* genes, as determined by RT-qPCR analysis, in RAF1/EGFR Resistant KPeFC;*Raf1*^L/L^;*Egfr*^L/L^ tumor cell lines expressing control shRNAs (open bars) or shRNAs against the respective genes (solid bars). The P value was obtained using paired two-tailed t test. ** P<0.01. Error bars indicate mean ± SEM. **E**, Western blot analysis of RAF1, EGFR and STAT5 expression levels in whole cell-extracts of a representative RAF1/EGFR resistant KPeFC;*Raf1*^L/L^;*Egfr*^L/L^ tumor cell line expressing non-targeted shRNAs (Control) or shRNAs against *Stat5* and infected with Adeno-GFP or Adeno-CRE viral particles. GAPDH was used as loading control.

**Figure S3.**
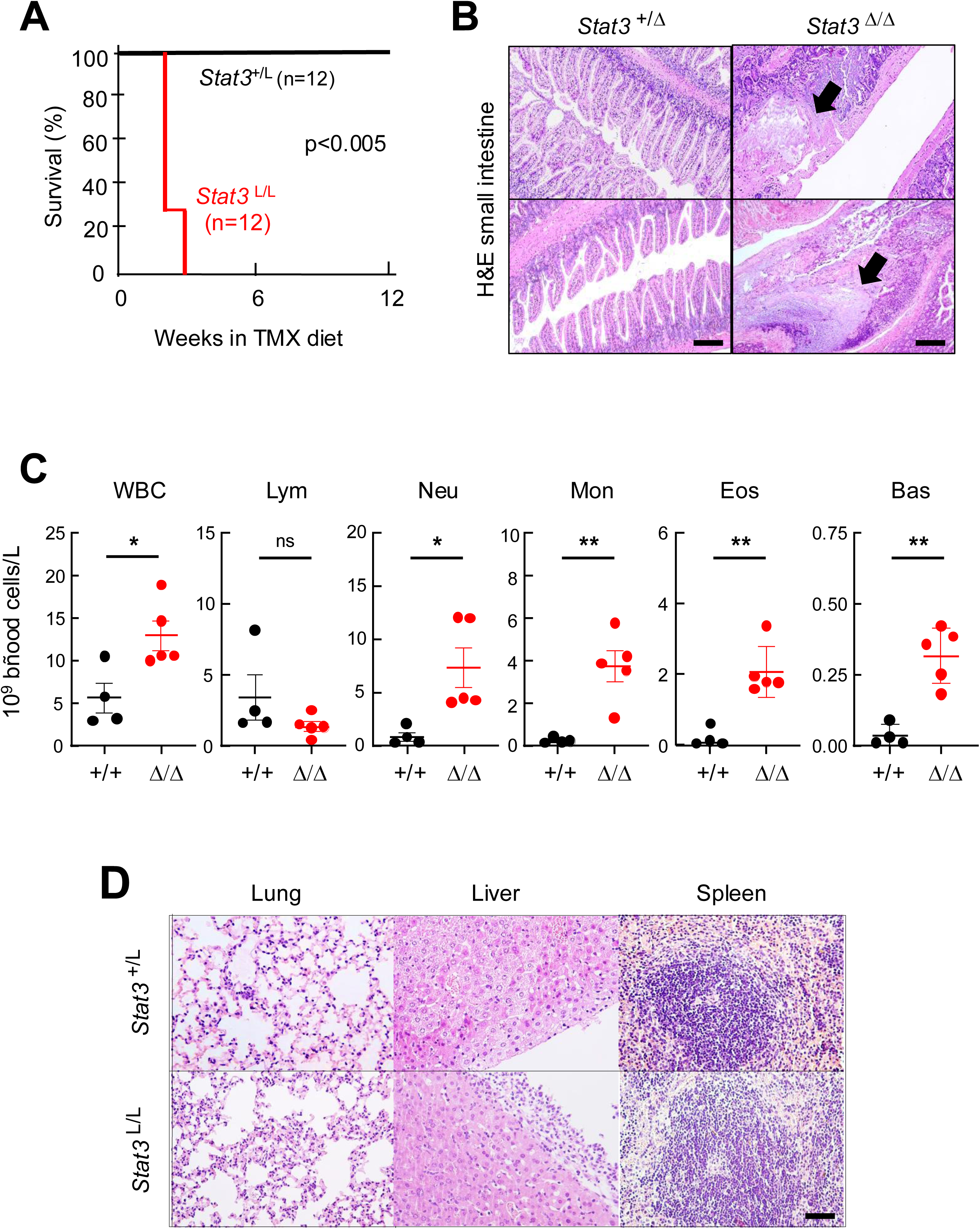
Differential toxicities induced by systemic genetic ablation versus systemic degradation of STAT3. **A**, Survival of adult (n=12) *Stat3*^+/L^;*Rosa26*-CreERT2 (solid line) and *Stat3*^L/L^;*Rosa26*-CreERT2 (red line) mice exposed to a TMX diet for 12 weeks. The P value was obtained using Log-rank Mantel-Cox test, p<0.005. **B**, Representative images of H&E-stained sections of the small intestine of *Stat3*^+/L^;*Rosa26*-CreERT2 and *Stat3*^L/L^; *Rosa26*-CreERT2 mice exposed to the TMX diet and sacrificed after 12 weeks of TMX exposure (*Stat3*^+/L^) or at humane endpoint (*Stat3*^L/L^), respectively. Solid arrows indicate representative intestinal ulcers. Scale bar represents 100 μm. **C**, Blood cell counts of white blood cells (WBC), lymphocytes (Lym), neutrophils (Neu), monocytes (Mon), eosinophils (Eos) and basophils (Bas) of wild type (+/+) (closed circles) (n=4) or *Stat3*^L/L^;*Rosa26*-CreERT2 (n=5) mice exposed to TMX diet for two weeks (Δ/Δ) (red circles). The P value was obtained using unpaired two-tailed t test. ns, not significant, * P<0.05, ** P<0.01. Error bars indicate mean ± SEM. **D**, H&E staining of representative images of sections of the indicated tissues of *Stat3*^+/L^;*Rosa26*-CreERT2 and *Stat3*^L/L^;*Rosa26*-CreERT2 mice (n=3) exposed to a TMX diet for two weeks. Scale bar represent 50 μm.

**Figure S4.**
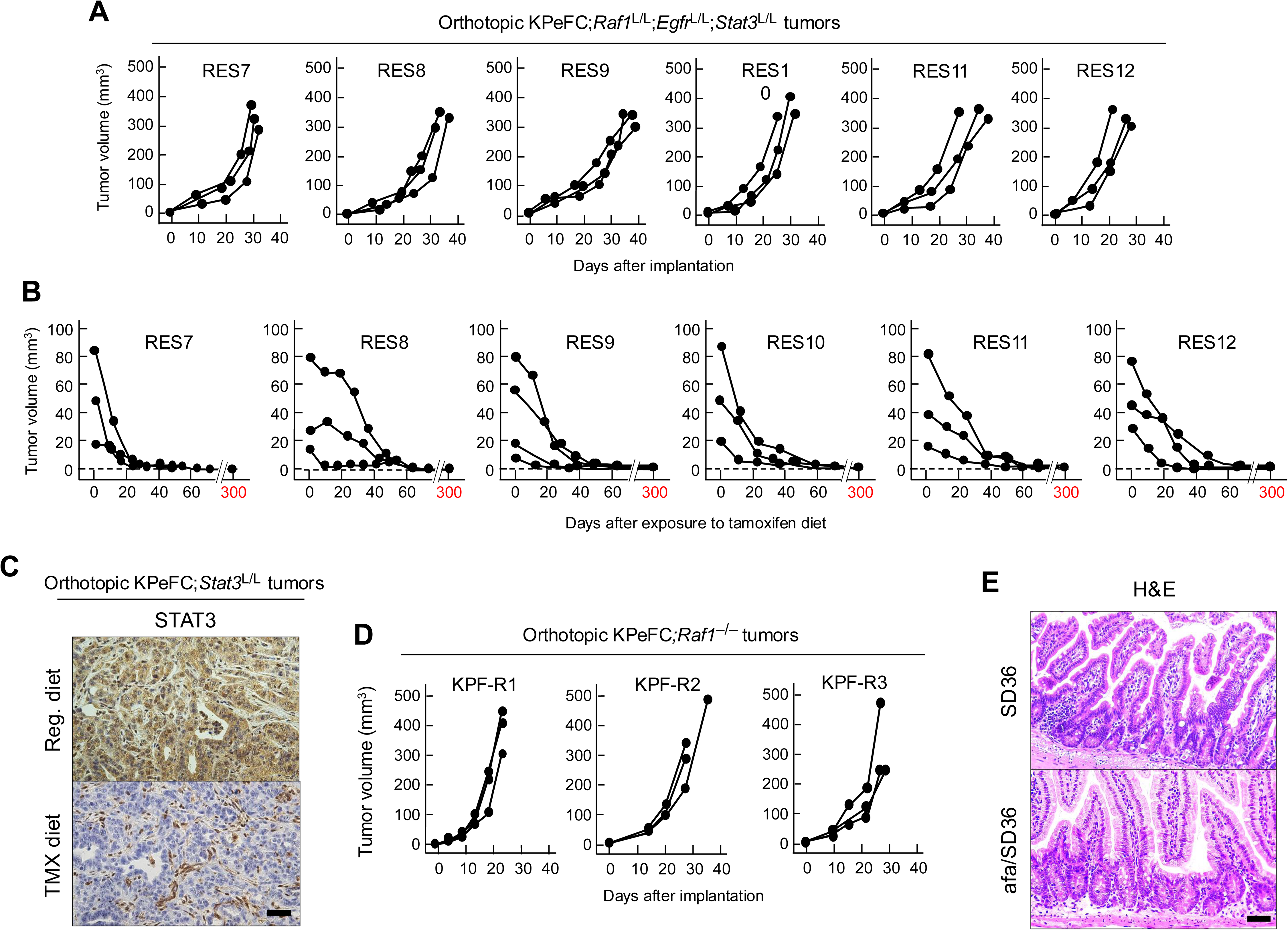
Therapeutic effect of concomitant ablation of *Raf*1, *Egf*r and *Stat3* in orthotopic pancreatic tumors. **A,** Tumor volume visualized by ultrasound imaging of tumors present in C57BL/6 mice (n=3) orthotopically implanted with six of twelve RAF1/EGFR resistant KPeFC; *Raf1*^L/L^;*Egfr*^L/L^*;Stat3*^L/L^ tumor cell lines (RES7 to RES12) fed with regular diet. **B**, Tumor volume visualized by ultrasound imaging of tumors present in C57BL/6 mice (n=3) orthotopically implanted with six of twelve RAF1/EGFR resistant KPeFC; *Raf1*^L/L^;*Egfr*^L/L^*;Stat3*^L/L^ tumor cell lines (RES7 to RES12) exposed to a TMX diet at the indicated times (red arrow) for 80 days. Mice were subsequently fed with regular diet until they reached 300 days port-implantation. Regular ultrasound analysis failed to revealed tumor relapse in any of the 18 mice. Dotted line represents tumor sizes undetectable by ultrasound. **C**, Representative IHC analysis of STAT3 expression in orthotopic KPeFC;*Stat3*^L/L^ tumors after 2 weeks of TMX exposure. Scale bar represents 100 μm **D**, Tumor volume visualized by ultrasound of orthotopic tumors of C57BL/6 mice (n=3) implanted with three independent KPeFC;*Raf1*^-/-^ tumor cell lines (KPF-R1 to KPF-R3) treated with vehicle when tumors reached 60 to 100 mm^3^ until the tumors reached humane endpoint. **E**, Representative images of H&E-stained sections of the intestinal epithelium of C57BL/6 mice implanted with a KPeF tumor cell line and treated for 8 weeks with SD36 (50 mpk, ip, qd) with or without afatinib (20 mpk, po, qd), respectively. Scale bar represents 100 μm.

**Figure S5.**
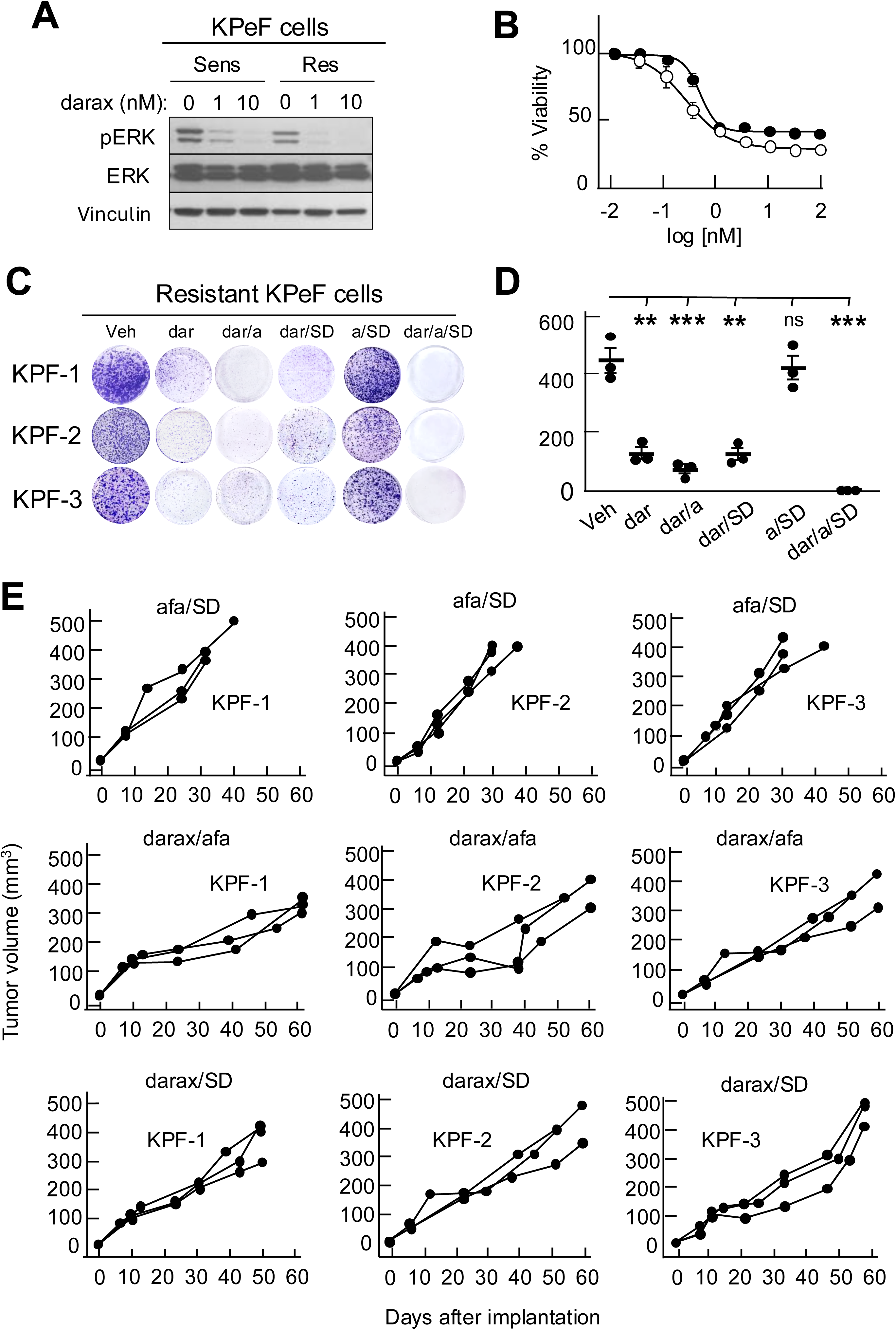
Dual combinations of inhibitors have no therapeutic activity in orthotopic pancreatic tumors. **A,** Western blot analysis of pERK and ERK expression in Sensitive and Resistant KPeF tumor cells upon treatment with the indicated concentrations of daraxonrasib for 24 h. Vinculin served as a loading control. **B**, Viability of three representative Sensitive (open circles) or Resistant (solid circles) KPeFC;*Raf1*^L/L^;*Egfr*^L/L^ tumor cell lines exposed to the indicated concentrations of daraxonrasib for 72 hours. Error bars represent mean ± SEM. **C**, Colony formation assay of three representative RAF1/EGFR resistant KPeF tumor cell lines (KPF-1 to KPF-3) treated with vehicle or with the indicated inhibitor, daraxonrasib (dar) (1 nM), afatinib (a) or SD36 (SD) (0.5 μM) for 10 days. **D**, Quantification of the number of colonies shown in **C**. The P value was obtained using multiple unpaired t tests. ns= not significant, **P<0.01 and ***P<0.001. Error bars indicate mean ± SEM. **E**, Tumor volume visualized by ultrasound of orthotopic tumors of C57BL/6 mice (n=3) implanted with three independent KPeF tumor cell lines (KPF-1 to KPF-3) treated with the indicated dual combinations of inhibitors (daraxonrasib (darax) (20 mpk, po, qd), afatinib (afa) (20 mpk, po, qd) and SD36 (SD) (50 mpk, ip, qd)) when the implanted tumors reached 120 to 200 mm^3^ in size for 60 days post-implantation or when the mice reached humane endpoint. All mice were eventually sacrificed when tumors reached humane endpoint.

**Figure S6.**
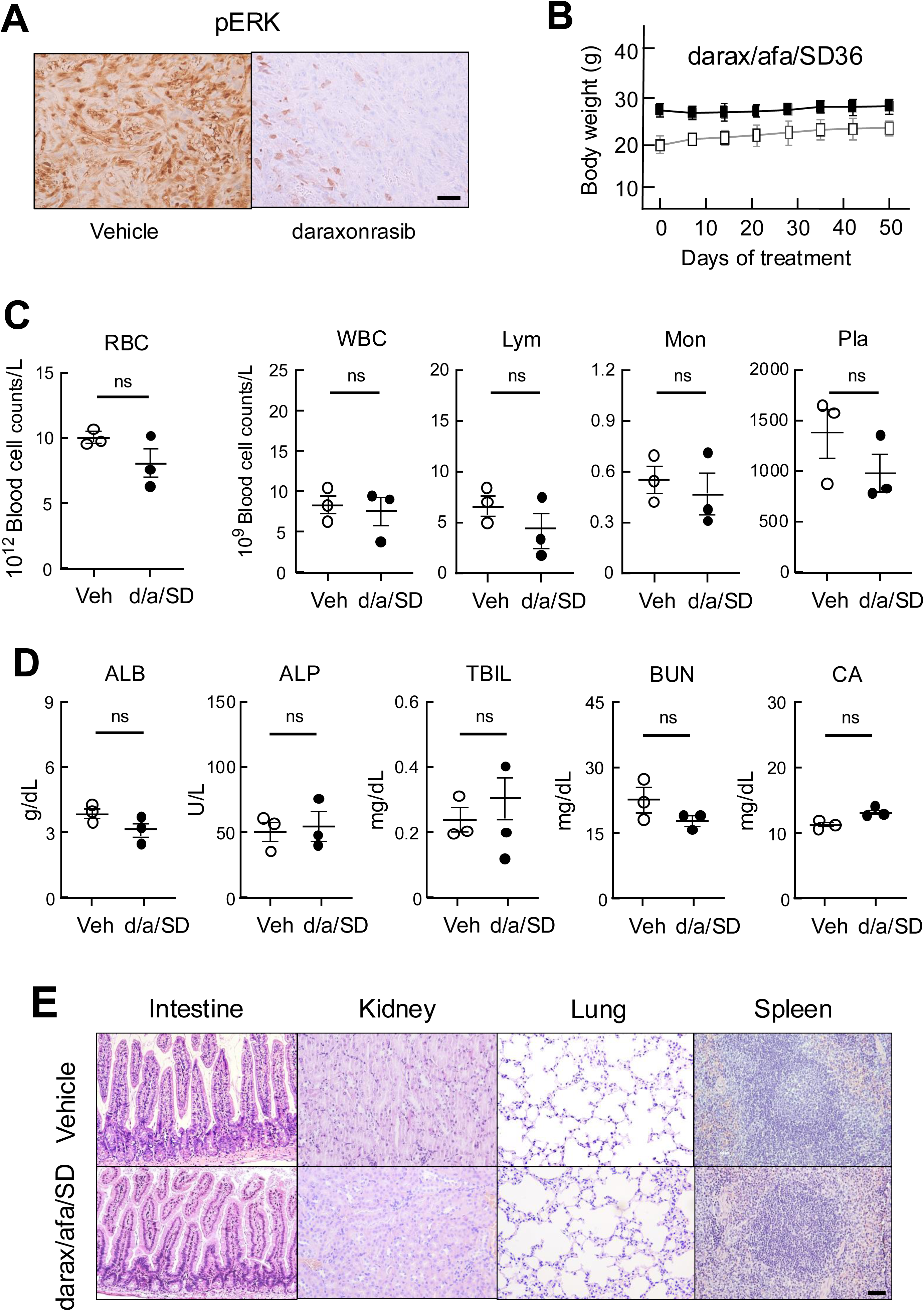
Tolerability of the triple daraxonrasib/afatinib/SD36 combination. **A,** Representative IHC analysis of pERK expression in orthotopic KPeF tumors treated with vehicle or daraxonrasib (20 mpk, po, qd) for four days. Mice (n=3) were sacrificed three hours after the treatment. **B**, Body weight of tumor-bearing C57BL/6 male (n=2) (solid squares) and female (n=4) (open squares) mice treated with a combination of daraxonrasib (20 mpk, po, qd), afatinib (20 mpk, po, qd) and SD36 (50 mpk, ip, qd) for the indicated periods of time. Error bars indicate mean ± SD. **C**, Blood cell counts of red blood cells (RBC), white blood cells (WBC), lymphocytes (Lym), monocytes (Mon) and platelets (Pla) of C57BL/6 mice treated with vehicle (open circles) (n=3) or with a combination of daraxonrasib (20 mpk, po, qd), afatinib (20 mpk, po, qd) and SD36 (50 mpk, ip, qd) (d/a/SD) (solid circles) (n=3) for three weeks. The P value was obtained using unpaired two-tailed t test. ns, not significant. Error bars indicate mean ± SEM. **D**, Metabolic parameters including albumin (ALB), alkaline phosphatase (ALP), total bilirubin (TBIL), blood urea nitrogen (BUN) and calcium (CA) of C57BL/6 mice treated with vehicle (open circles) (n=3) or with a combination of daraxonrasib (20 mpk, po, qd), afatinib (20 mpk, po, qd) and SD36 (50 mpk, ip, qd) (d/a/SD) (solid circles) (n=3) for three weeks. The P value was obtained using unpaired two-tailed t test. ns, not significant. Error bars indicate mean ± SEM. **E**, H&E staining of representative images of sections of the indicated tissues obtained from tumor-bearing orthotopic KPeF mice (n=3) exposed to (**top**) vehicle or to (**bottom**) a combination of daraxonrasib (20 mpk, po, qd), afatinib (20 mpk, po, qd) and SD36 (50 mpk, ip, qd) (darax/afa/SD) for three to eight weeks. Scale bar represent 50 μm.

**Figure S7.**
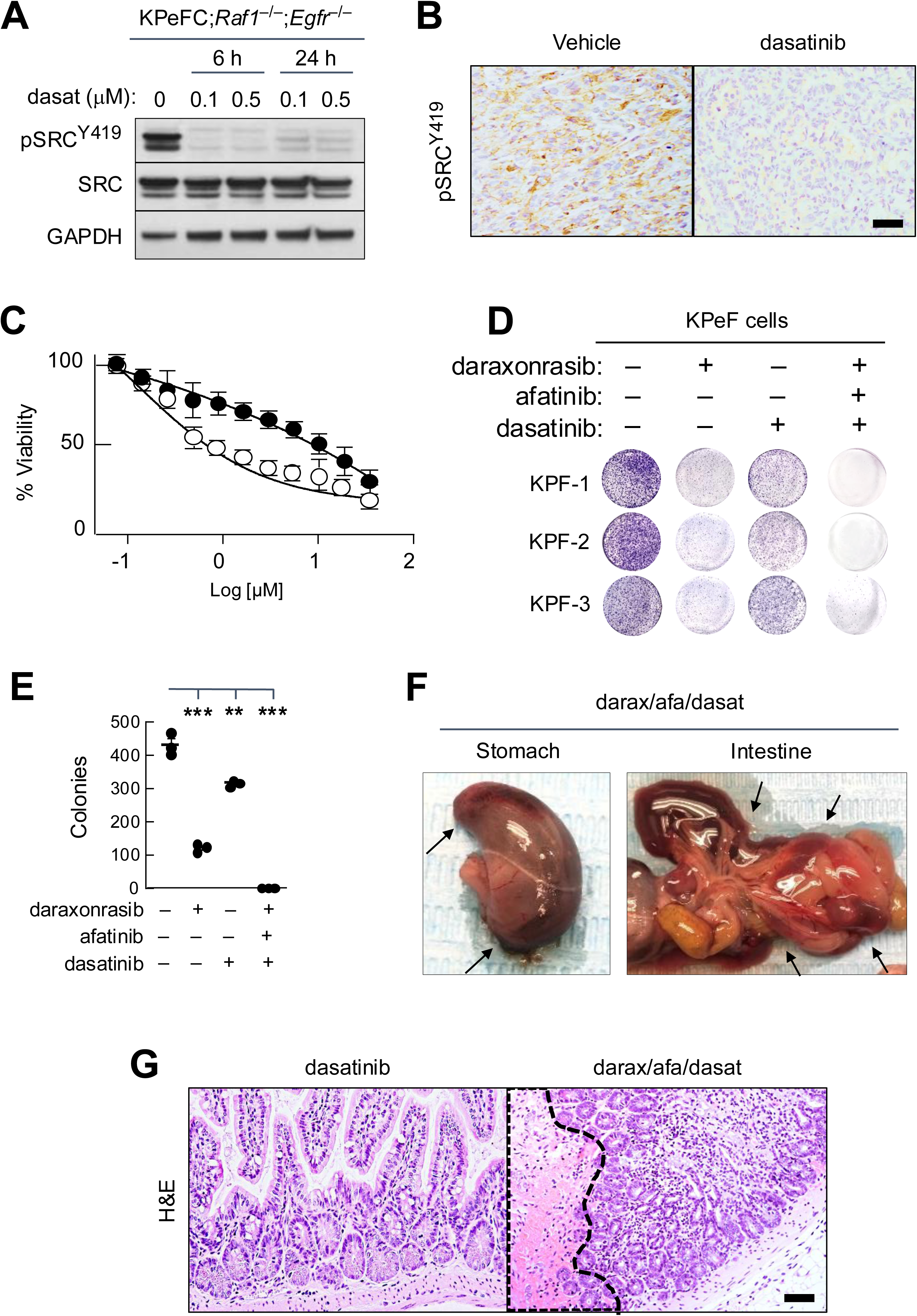
Mice treated with a combination of daraxonrasib/afatinib/dasatinib develop gastric and intestinal hemorrhages. **A,** Western blot analysis of pSRC^Y419^ and SRC expression in a representative KPeFC;*Raf1*^-/-^;*Egfr*^-/-^ tumor cell line treated with the indicated concentrations of dasatinib for 6 or 24 h. GAPDH was used as loading control. **B**, Representative IHC analysis of pSRC^Y419^ expression in sections of an orthotopic KPeFC;*Raf1*^-/-^;*Egfr*^-/-^ tumor treated with either Vehicle or dasatinib (5 mpk, po, qd) for three days and sacrificed three hours after treatment. Scale bar represent 50 mm. **C**, Viability of three representative RAF1/EGFR resistant KPeFC;*Raf1*^L/L^;*Egfr*^L/L^ (solid circles) and KPeFC;*Raf1*^-/-^;*Egfr*^-/-^ (open circles) tumor cell lines exposed to the indicated concentrations of dasatinib for 72 hours. Error bars represent mean ± SEM. **D**, Colony formation assay of three representative RAF1/EGFR resistant KPeF tumor cell lines (KPF-1 to KPF-3) untreated or treated with daraxonrasib (1 nM), afatinib (0.5 μM) and dasatinib (0.5 μM) for 10 days. **E**, Quantification of the number of colonies shown in **D**. The P value was obtained using multiple unpaired t tests **P<0.01 and ***P<0.001. Error bars indicate mean ± SEM. **F**, A representative images of gastric and intestinal tissue obtained during the necropsy of mice treated with daraxonrasib (20 mpk, po, qd), afatinib (20 mpk, po, qd) and dasatinib (5 mpk, ip, qd) (darax/afa/dasat). Mice were sacrificed at human endpoint within the first 24 h of treatment. Arrows indicate areas of hemorrhage. **G**, H&E analysis of the intestine of mice treated either with dasatinib (5 mpk, ip, qd) for one week or with the indicared triple combination within the first 24 h of treatment. Dotted line encircles an area of hemorrhage. Scale bar represents 50 μm.

**Figure S8.**
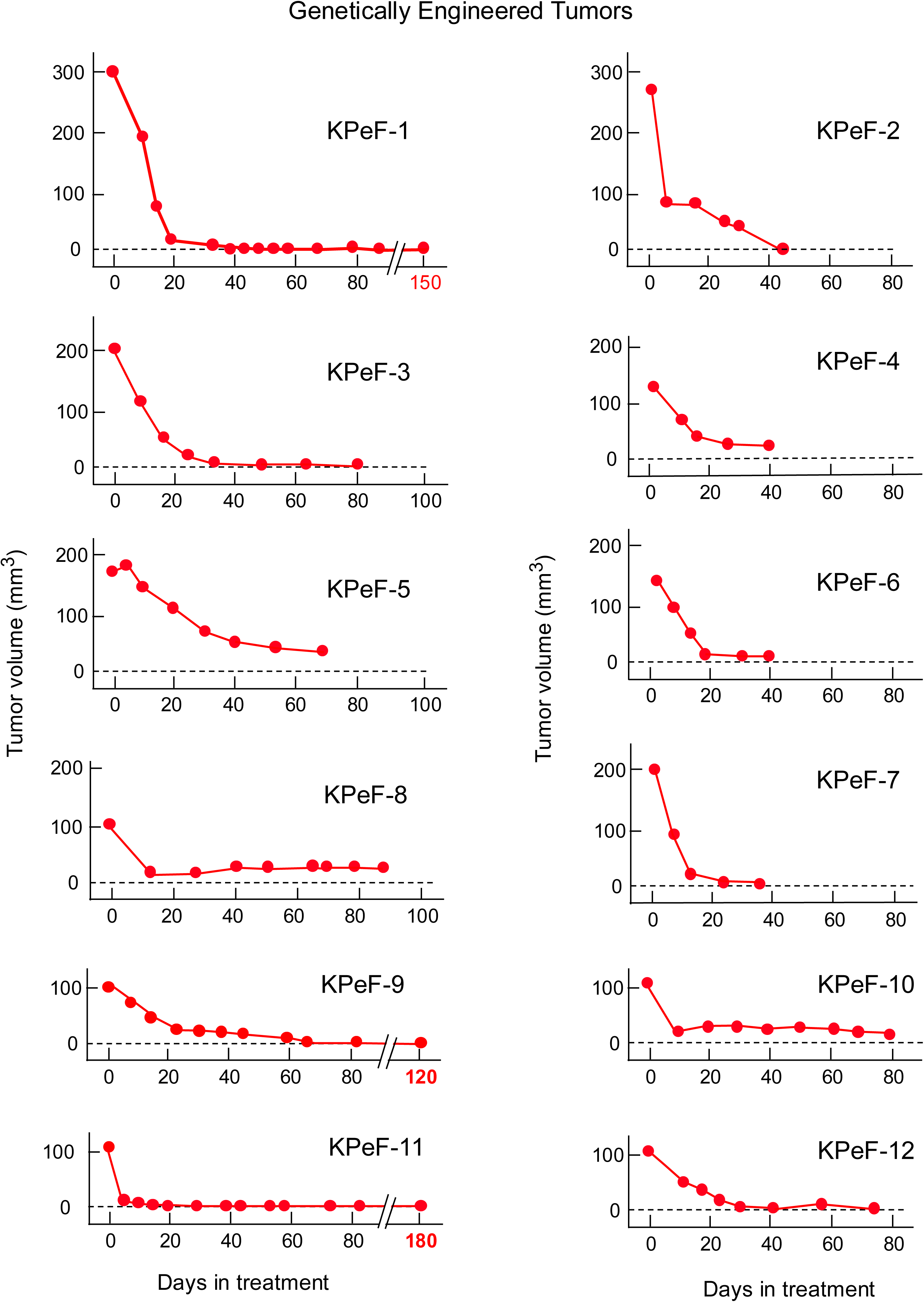
Therapeutic effect of the triple combination in genetically engineered mouse tumors. Individual representation of genetically engineered tumors developed by KPeF mice (KPeF-1 to KPeF-12) treated with a combination of daraxonrasib (20 mpk, po, qd), afatinib (20 mpk, po, qd) and SD36 (50 mpk, ip, qd). Treatment started when tumors reached at least 100 mm^3^ (0 time points). Dotted line represents tumor sizes undetectable by ultrasound.

**Figure S9.**
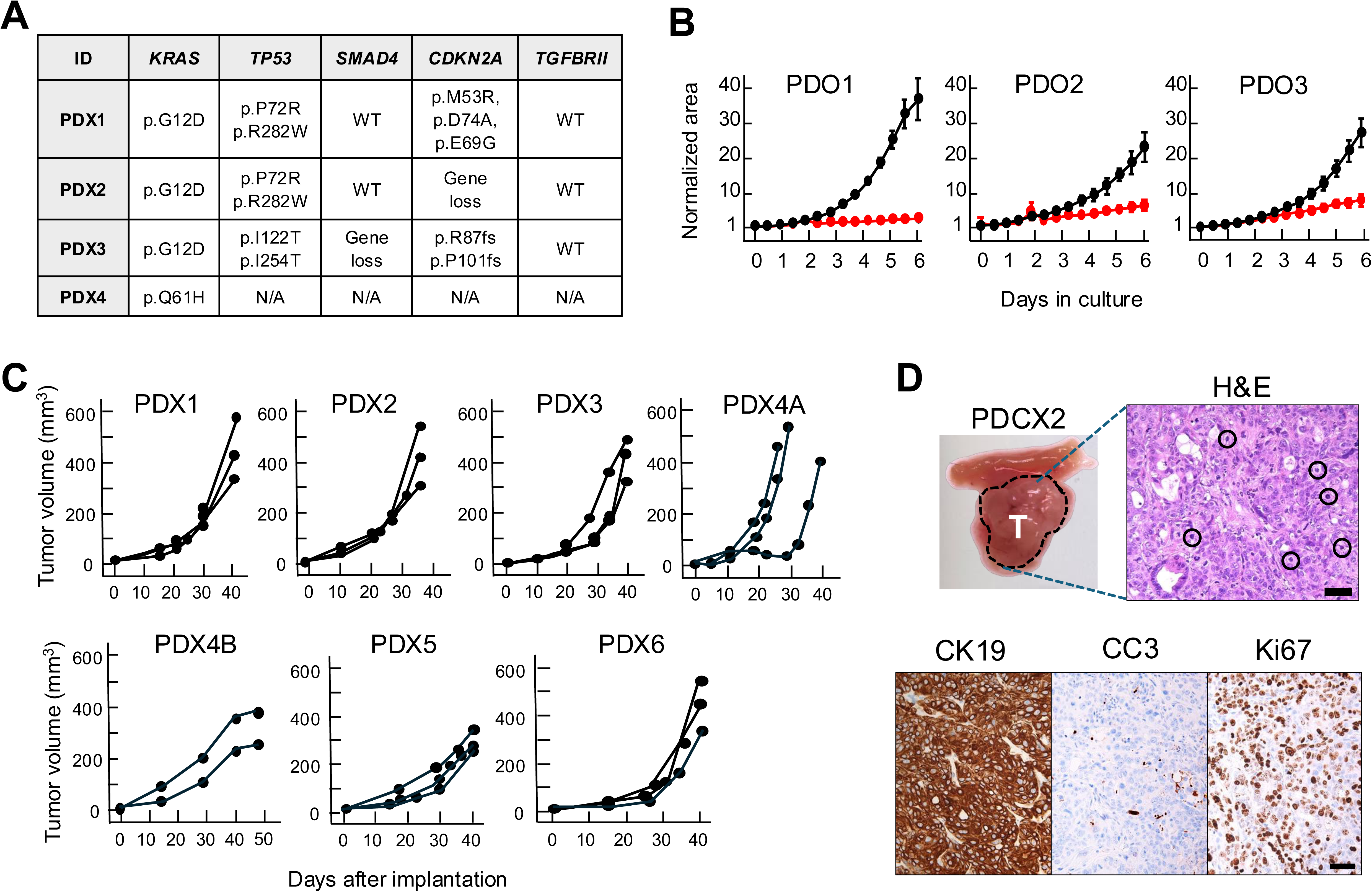
PDO and PDX tumor models: Mutational spectra and tumor growth. **A,** Mutational status of KRAS, TP53, SMAD4, CDKN2A and TGFBRII in the indicated PDX tumor models, N/A Not available. **B**, Normalized area occupied by PDO cultures treated with vehicle (solid circles) or with the combination of KRas^G12D^ inhibitor MRTX1133 (0.5 μM), afatinib (0.5 μM) and SD36 (1 μM) (red circles). **C**, Tumor volume of PDX1, PDX2, PDX3 and PDX4A, PDX-4B, PDX5 and PDX6-derived tumors implanted in immunodeficient mice (n=3), visualized by ultrasound. Mice were treated with vehicle when tumors reached 100 to 120 mm^3^. Mice were sacrificed at humane endpoint when they reached 400 to 600 mm^3^. **D**, (**Top**) Representative image of a pancreatic tumor (T) isolated from a nude mouse orthotopically implanted with the PDX2 tumor model exposed to vehicle. Representative mitotic figures in an H&E section are indicated by circles. (**Bottom**) Images of Cytokeratin 19 (CK19) Cleaved Caspase 3 (CC3) and Ki67 stained sections of the pancreatic tumor shown above. Scale bar represents 50 μm.

**Figure S10.**
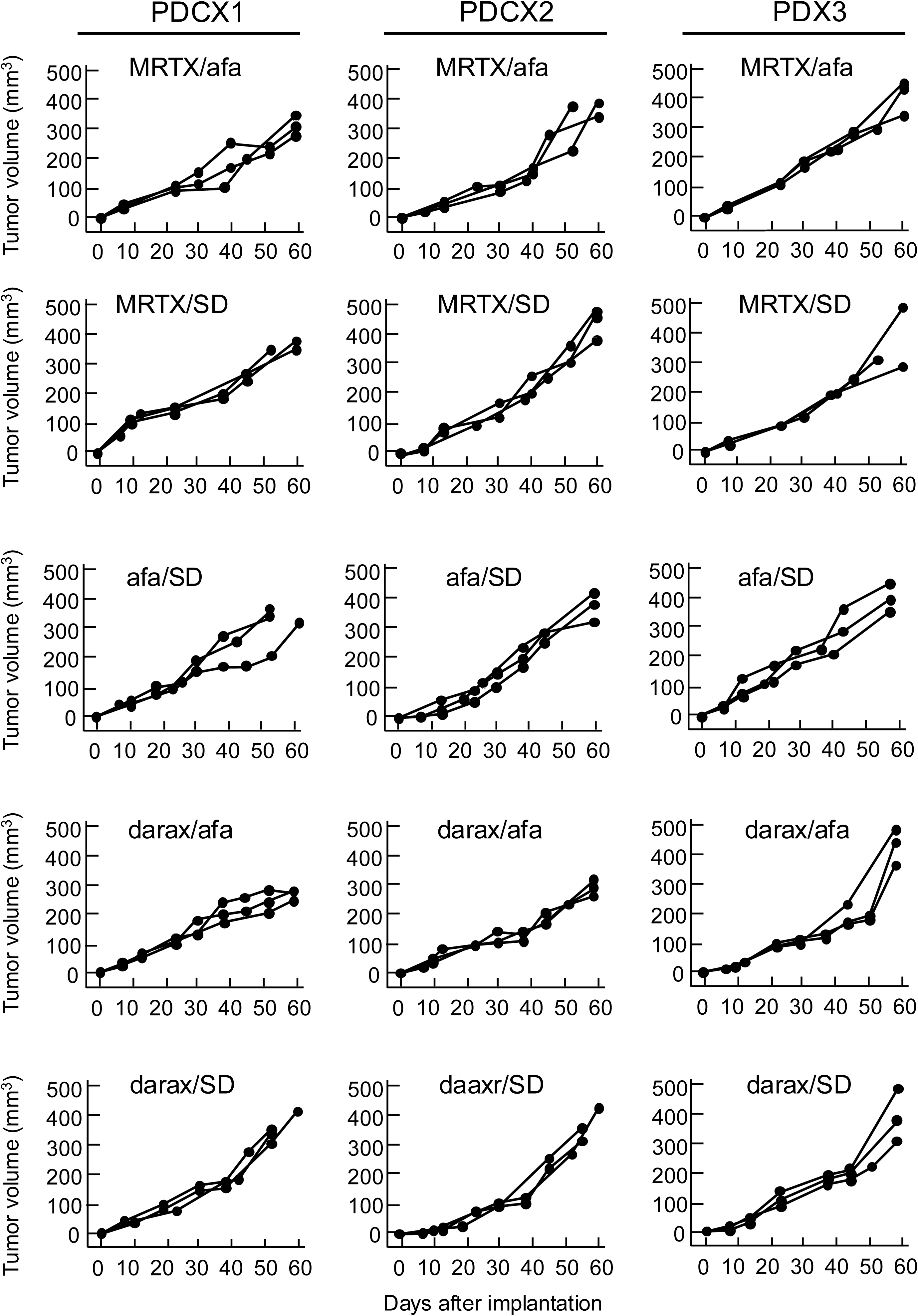
Lack of therapeutic activity of dual drug combinations in PDX models. Tumor volume of PDX1 and PDX2 and PDX3 tumor models orthotopically implanted in immunodeficient mice (n=3) and visualized by ultrasound. Mice were treated with dual combinations of daraxonrasib (20 mpk, po, qd), MRTX1133 (30 mpk, ip, qd), afatinib (20 mpk, po, qd) and SD36 (50 mpk, ip, qd) when tumors reached 100 to 120 mm^3^. Mice were sacrificed at humane endpoint when they reached 300 to 500 mm^3^.

**Figure S11.**
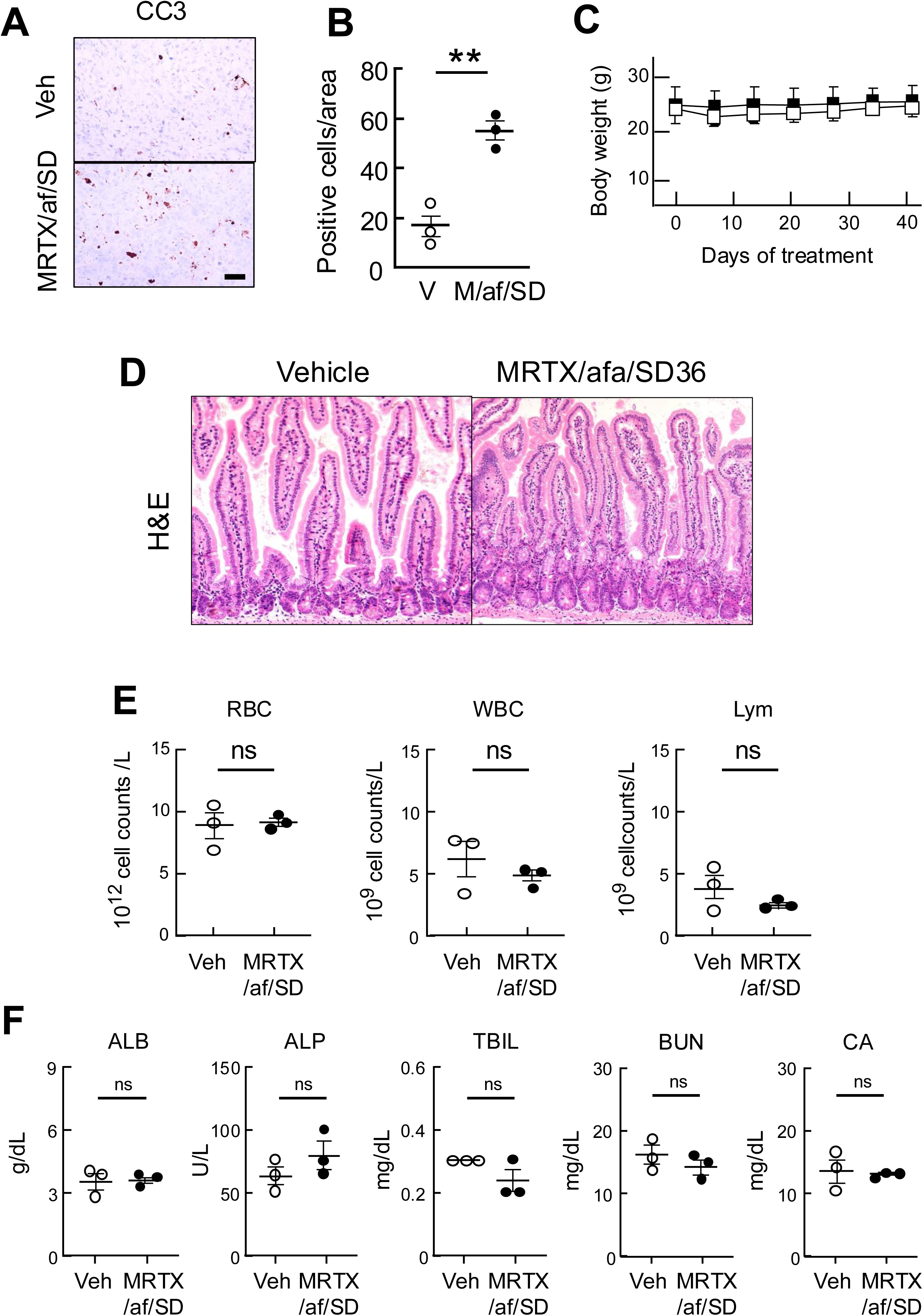
Mechanisms of PDX tumor regression and lack of toxicity of the triple combination of MRTX1133/aftinib/SD36 in immunocompromised mice. **A**, IHC analysis of CC3 expression in tumors of around 100 mm^3^ (n=3) exposed for four days to vehicle (Veh) or to the triple combination, MRTX1133 (30 mpk, ip, qd), afatinib (20 mpk, po, qd) and SD36 (50 mpk, ip, qd) (MRTX/a/SD) and sacrificed three hours after the treatment. Scale bar represents 50 μm. **B**, Quantification of the number of CC3 positive cells per area depicted in **A**. The results represent the average of CC3 in three independent tumors treated with vehicle (open circles) or with the triple combination (M/af/SD) (solid circles). The P value was obtained using unpaired two-tailed t test. **P<0.01. Error bars indicate mean ± SEM. **C**, Body weight of tumor-bearing nude female mice (n=5) treated with vehicle (open squares) or with a combination of MRTX1133 (30 mpk, po, qd), afatinib (20 mpk, po, qd) and SD36 (50 mpk, ip, qd) (solid squares) for the indicated periods of time. Error bars indicate mean ± SD. **D,** Representative images of H&E stained sections of the intestinal epithelium of a nude mouse implanted with the PDX2 tumor model and treated with either vehicle or a combination of MRTX1133 (30 mpk, po, qd), afatinib (20 mpk, po, qd) and SD36 (50 mpk, ip, qd) (MRTX/afa/SD36). Scale bar represents 100 μm. **E**, Blood cell counts of red blood cells (RBC), white blood cells (WBC), and lymphocytes (Lym) of immunocompromised mice treated with vehicle (open circles) (n=3) or with a combination of MRTX1133 (30 mpk, ip, qd), afatinib (20 mpk, po, qd) and SD36 (50 mpk, ip, qd) (MRTX/af/SD) (solid circles) (n=3) for three weeks. The P value was obtained using unpaired two-tailed t test. ns, not significant. Error bars indicate mean ± SEM. **F**, Metabolic parameters including albumin (ALB), alkaline phosphatase (ALP), total bilirubin (TBIL), blood urea nitrogen (BUN) and calcium (CA) of immunocompromised mice treated with vehicle (open circles) (n=3) or with a combination of MRTX1133 (30 mpk, ip, qd), afatinib (20 mpk, po, qd) and SD36 (50 mpk, ip, qd) (MRTX/af/SD) (solid circles) (n=3) for three weeks. The P value was obtained using unpaired two-tailed t test. ns, not significant. Error bars indicate mean ± SEM.

